# Facultative chemosynthesis in a deep-sea anemone from hydrothermal vents in the Pescadero Basin, Gulf of California

**DOI:** 10.1101/2020.08.10.245456

**Authors:** Shana K Goffredi, Cambrie Motooka, David A. Fike, Luciana C Gusmão, Ekin Tilic, Greg W Rouse, Estefanía Rodríguez

## Abstract

**Background:** Numerous deep-sea invertebrates have formed symbiotic associations with internal chemosynthetic bacteria in order to harness inorganic energy sources typically unavailable to most animals. Despite success in nearly all marine habitats and their well-known associations with photosynthetic symbionts, Cnidaria remain one of the only phyla without a clear dependence on hydrothermal vents and reliance on chemosynthetic bacterial symbionts specifically.

**Results:** A new chemosynthetic symbiosis between the sea anemone *Ostiactis pearseae* (Daly & Gusmão, 2007) and intracellular bacteria was discovered at ~3700 m deep hydrothermal vents in the southern Pescadero Basin, Gulf of California. Unlike most sea anemones observed from chemically-reduced habitats, this species was observed in and amongst vigorously venting fluids, side-by-side with the chemosynthetic tubeworm *Oasisia* aff. *alvinae.* Individuals of *O. pearseae* displayed carbon, nitrogen, and sulfur tissue isotope values (average δ^13^C −29.1‰, δ^15^N 1.6‰, and δ^34^S −1.1‰) suggestive of a distinct nutritional strategy from conventional Actiniaria suspension feeding or prey capture. Molecular and microscopic evidence confirmed the presence of intracellular SUP05-related bacteria housed in the tentacle epidermis of *O. pearseae* specimens collected from 5 hydrothermally-active structures within two vent fields ~2 km apart. SUP05 bacteria dominated the *O. pearseae* bacterial community (64-96% of the total bacterial community based on 16S rRNA sequencing), but were not recovered from other nearby anemones, and were generally rare in the surrounding water (< 7% of the total community). Further, the specific *Ostiactis*-associated SUP05 phylotypes were not detected in the environment, indicating a specific association. Two unusual candidate bacterial phyla (the OD1 and BD1-5 groups) also appeared to associate exclusively with *O. pearseae* and may play a role in symbiont sulfur cycling.

**Conclusion:** *Ostiactis pearseae* represents the first member of Cnidaria described to date to have a physical and nutritional alliance with chemosynthetic bacteria. The facultative nature of this symbiosis is consistent with the dynamic relationships formed by both the SUP05 bacterial group and Anthozoa. The advantages gained by appropriating metabolic and structural resources from each other presumably contribute to their striking abundance in the Pescadero Basin, at the deepest known hydrothermal vents in the Pacific Ocean.

## Background

Numerous deep-sea annelids, molluscs, and other invertebrates have forged relationships with bacteria in order to harness inorganic sources of energy that are typically unavailable to most animals. Microbial chemosynthesis generates energy through the oxidation of sulfide, as an example, used to fuel the production of organic carbon, which can be shared with a receptive animal host. To date, members of at least six major animal clades, including most recently *Trichoplax* (Placozoa), have formed symbiotic associations with internal chemosynthetic bacteria (Dubilier 2008; Gruber-Vodicka et al. 2019). Interestingly, Cnidaria, although well-known to host photosynthetic symbionts, remains one of the last prominent animal clades without a documented metabolic dependence on chemosynthetic bacterial symbionts for survival in hydrothermal vents.

Anthozoa, which includes sea anemones and corals, is among the most successful and diverse group of Cnidaria, found in all marine habitats at most depths and latitudes (Daly et al. 2008). Their worldwide ecological success may best be attributed to an ability to form symbiotic relationships with other organisms, including microbial eukaryotes (e.g. the dinoflagellate *Symbiodinium*; LaJeunesse et al. 2018), as in the case of shallow-water tropical species. Anthozoa such as sea anemones, octocorals and zoanthids are also found at deep-sea reducing environments, such as hydrothermal vents, seeps, and whalefalls (Daly and Gusmão 2007; Zelnio et al 2009; Rodríguez et al. 2012; Breedy et al. 2019), however they have been historically understudied, and most remain undescribed. It would be reasonable, and perhaps even expected, for some of these deep-sea Anthozoa to also host microbial symbionts. In fact, a recent study demonstrated an affiliation between sulfide-oxidizing bacteria and certain species of Anthozoa found near deep-sea methane seeps, however the specific mode and importance of this relationship is not yet known (Vohsen et al. 2020).

The recently discovered Pescadero Basin vent field at 3700 m depth in the southern Gulf of California differs markedly from nearby vent localities (e.g. Guaymas Basin and 21°N East Pacific Rise) in physical, chemical, and biological attributes (Caress et al. 2015; Goffredi et al. 2017; Paduan et al. 2018). In particular, the vents in the Pescadero Basin are uniquely composed of hydrothermal calcite, with venting fluids that contain high levels of aromatic hydrocarbons, hydrogen, methane and hydrogen sulfide at a pH of ~6.5 (Goffredi et al. 2017). The Pescadero Basin vents are also highly unusual in faunal composition with many new species and numerous others that do not occupy nearby regional vents (e.g. Alarcon Rise vents; Rouse et al. 2016; Goffredi et al. 2017; Hatch et al. 2020). Included in this group of unusual fauna was a very abundant white sea anemone (up to 68 individuals m^-2^ in some areas) that occurred in and amongst the siboglinid tubeworm *Oasisia* aff. *alvinae*, often very near to actively venting fluids (Fig. 1; Supplemental Video; Goffredi et al. 2017).

**Figure 1:**
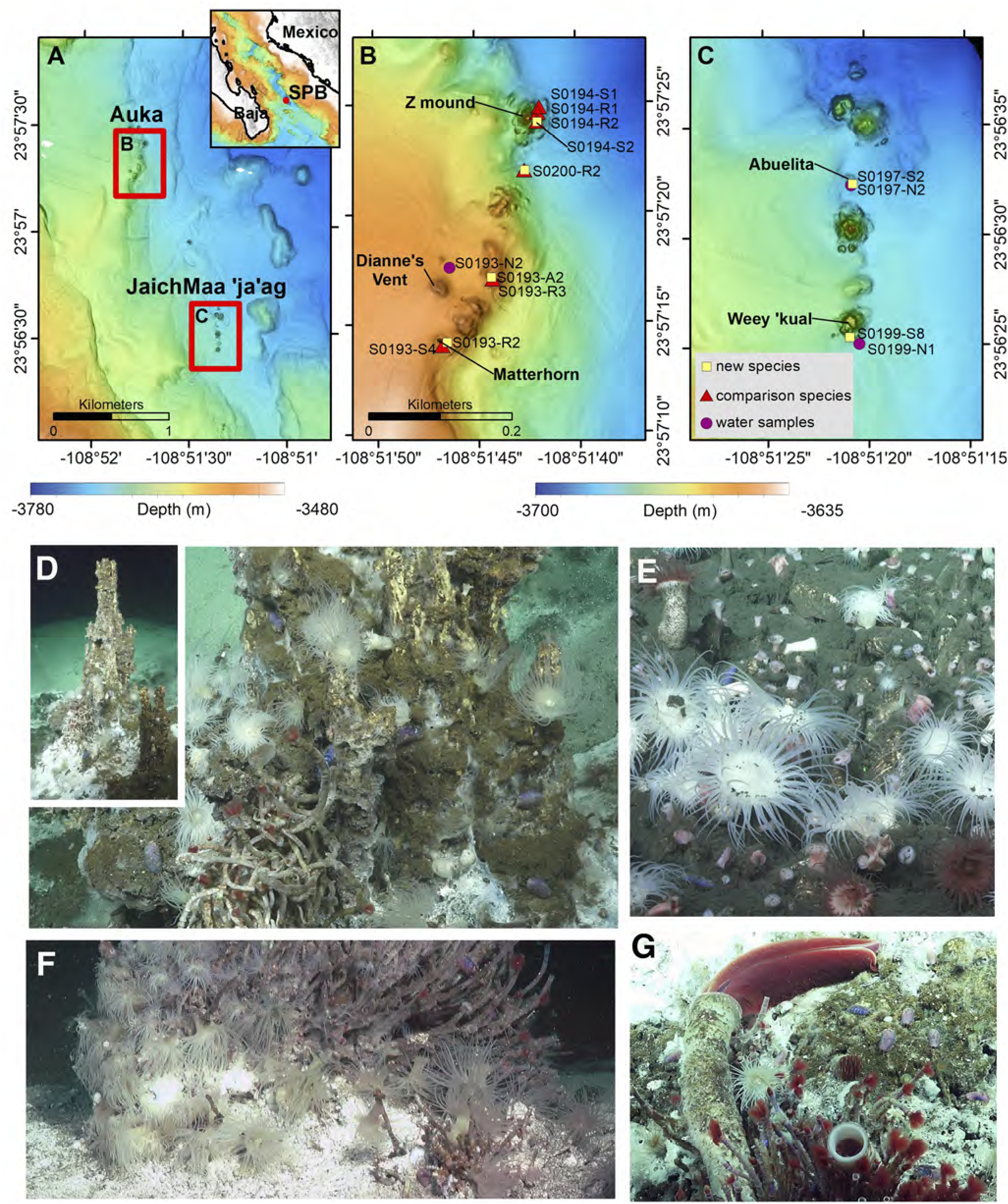
Locations and in situ images of the actiniarian *Ostiactis pearseae*. **A.** Location of South Pescadero Basin (SPB) vent fields Auka (in B) and JaichMaa ‘ja’ag (in C). Inset shows location of SPB at the mouth of the Gulf of California between the tip of the Baja Peninsula and mainland Mexico. **B.** Auka vent field samples and chimneys. (samples symbolized as in C). **C.** JaichMaa ‘ja’ag vent field samples and chimneys. Legend shows the sample types. Maps A,B, and C show 1-m resolution bathymetry collected by mapping AUVs (owned and operated by the Monterey Bay Aquarium Research Institute). Color ramps show the depth ranges. **D-F.** Specimens of *Ostiactis pearseae* collected from both vent fields (shown as yellow squares in B and C), indicated by arrows. **G.** An individual *O. pearseae* near to the chemosynthetic tubeworms *Riftia pachyptila* and *Oasisia* aff*. alvinae.*

Previously, several unidentified Pescadero Basin Actiniaria (sea anemones) were reported to be quite depleted in tissue δ^13^C values (−33 to −38‰; Goffredi et al. 2017; Salcedo et al. 2019). This evidence, along with their unusual life position and abundance in zones of active fluid venting, hinted at their possible nutritional reliance on chemoautotrophic carbon production, as opposed to traditional suspension feeding or prey capture via cnidae, however, the specific details were not explored further. Here, by combining microbial community profiling, ultrastructural analysis via microscopy, and stable isotope measurements, we document the first species of chemosynthetic sea anemone at vents deep in the Gulf of California, identified as *Ostiactis pearseae* (previously known only from whalefalls; Daly and Gusmão 2007). This extensive new population of *O. pearseae* appears to rely on nutritional supplementation of carbon, nitrogen, and sulfur by intracellular bacteria within the SUP05 clade, housed in their epidermis.

## Results

Actiniaria of various morphotypes were observed to be one of only a handful of dominant animal species in both the Pescadero Basin Auka vent field (Goffredi et al. 2017; Paduan et al. 2018) and the newly discovered JaichMaa ‘ja’ag vent field, both within ~2 km of each other in the Gulf of California (Fig. 1). A conspicuous white actiniarian species visually represented a significant fraction of the animal community and was collected from 5 vent edifices in zones of active venting, very near to the obligate vent tubeworm, *Oasisia* aff. *alvinae* (Fig. 1). Several other sea anemones (by morphotype) were observed and collected near these same sites, usually in areas of less active fluid flow (Table 1). The white actiniarian morphotype was identified as *Ostiactis pearseae* (Daly and Gusmão, 2007), based on anatomical, cnidae and DNA sequencing of preserved polyps. The Pescadero Basin populations of *O. pearseae* showed slight differences in morphology and cnidae to the description of specimens from the type locality and, thus, an amendment to the species diagnosis is provided.

**Table 1:** Sample locations within the Pescadero Basin, Gulf of California, along with dive information, depth, and specimen descriptions for Anthozoa other than *Ostiactis pearseae.*

| *Ostiactis pearseae* |  |  |  |  |
| --- | --- | --- | --- | --- |
| Auka vent field | Dive #-Sample # | Depth | Lat | Lon |
| Matterhorn<sup>1</sup> | S0193-R2 | -3655 | 23°57.24218 N | 108°51.77758 W |
| E. of Diane's vent | S0193-A2 | -3655 | 23°57.28307 N | 108°51.73963 W |
| Z vent (top)<sup>1</sup> | S0194-S2 | -3670 | 23°57.39937 N | 108°51.70276 W |
| S. of Z vent (small chimney) | S0200-R2w | -3687 | 23°57.36546 N | 108°51.71477 W |
| JaichMaa 'ja'ag vent field |  |  |  |  |
| Abuelita<sup>1</sup> | S0197-S2 | -3692 | 23°56.53971 N | 108°51.34850 W |
| Weey 'kual | S0199-S8 | -3674 | 23°56.42347 N | 108°51.35257 W |
| Water samples<sup>2</sup> |  |  |  |  |
| Auka vent field | Dive #-Sample # | Depth | Lat | Lon |
| E. of Diane's vent | S0193-N2 | -3642 | 23°57.29596 N | 108°51.77752 W |
| JaichMaa 'ja'ag vent field |  |  |  |  |
| Abuelita | S0197-N2 | -3693 | 23°56.53998 N | 108°51.35074 W |
| Weey 'kual | S0199-N1 | -3669 | 23°56.41838 N | 108°51.34473 W |
| Other Anthozoa<sup>3</sup> |  |  |  |  |
| Auka vent field | Dive #-Sample # | Depth | Description |  |
| Matterhorn | S0193-S4 | -3655 | Kadosactinidae 'sp.B' Green tentacles |  |
| E. of Diane's vent | S0193-R3 | -3655 | Unidentified zoanthid |  |
| Z vent (lower on structure) | S0194-R1 | -3670 | Unidentified zoanthid |  |
| NW. of Z vent (diffuse flow) | S0194-R2 | -3687 | Unidentified zoanthid |  |
| NW. of Z vent (diffuse flow) | S0194-S1 | -3687 | Unidentified zoanthid |  |
| S. of Z vent (small chimney) | S0200-R2r | -3692 | Kadosactinidae 'sp.B' Red tentacles |  |

<sup>1</sup> anemones collected very near active *Oasisia* tubeworms
<sup>2</sup> collected via Niskin sampler aboard the ROV SuBastian
<sup>3</sup> collected in the same general venting structure as *O. pearseae*, geo-locations noted above

Class Anthozoa Ehrenberg, 1834

Subclass Hexacorallia Haeckel, 1896

Order Actiniaria Hertwig, 1882

Suborder Enthemonae Rodríguez and Daly, 2014 in Rodríguez et al. 2014

Superfamily Metridioidea Carlgren, 1893

Family Ostiactinidae Rodríguez et al. 2012

Genus *Ostiactis* Rodríguez et al. 2012

*Ostiactis pearseae* (Daly & Gusmão, 2007)

(Figures 2-3, Table 2; Table S2)

**Figure 2.**
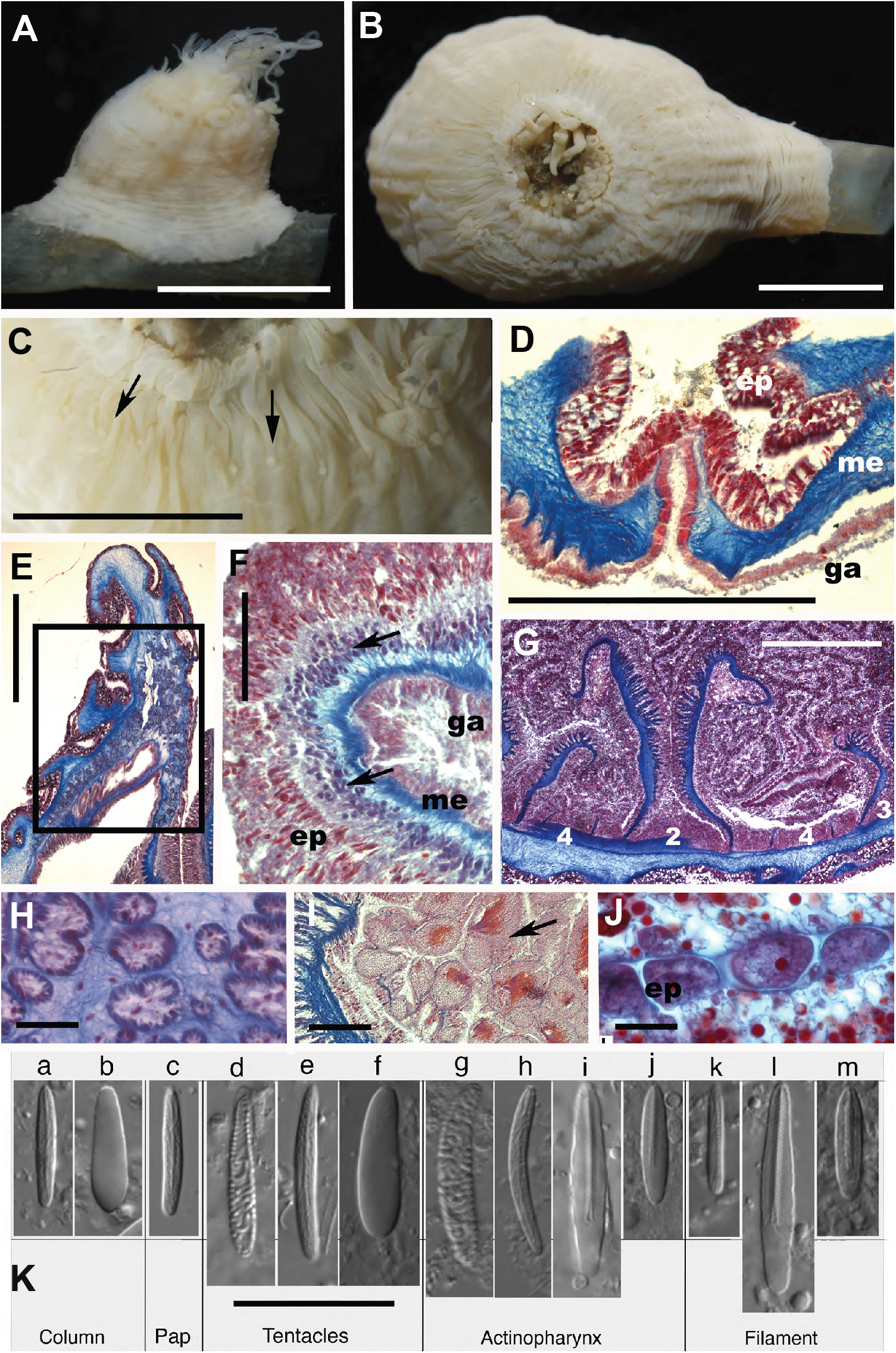
*Ostiactis pearseae* external and internal anatomy, including cnidae. **A-B.** External anatomy of *Ostiactis pearseae* from Pescadero Basin; (A) lateral view; (B) oral view. **C.** Detail of distal row of papillae in the column (arrows). **D.** Detail of longitudinal section through perforated papillae (cinclide). **E.** Longitudinal section of distal column showing mesogleal marginal sphincter muscle (area within rectangle). **F.** Cross section of a tentacle showing ectodermal longitudinal muscles (arrows). **G.** Cross section at the actinopharynx level showing cycles of mesenteries; numbers between mesenteries indicate different cycles. **H.** Detail of marginal sphincter muscle fibers in the mesoglea. **J.** Detail of developing oocytes and lipid inclusions (red small dots) in the gastrovascular cavity. **I.** Detail of spermatic cysts (arrow points to largest cyst). **K.** Cnidae types of *O. pearseae:* basitrichs (a, c, e, h, k), holotrichs (b, f), robust spirocysts (d, g), *p*-mastigophores A (i, l), *p*-mastigophores B1 (j, m). Abbreviations: ep, epidermis; ga, gastrodermis; me, mesoglea; pap, papillae. Scale bars: A-C, 6 mm; D, G, 1 mm; E, 0.5 mm; F, H, I, J, 0.1 mm; K, 25 μm.

**Figure 3.**
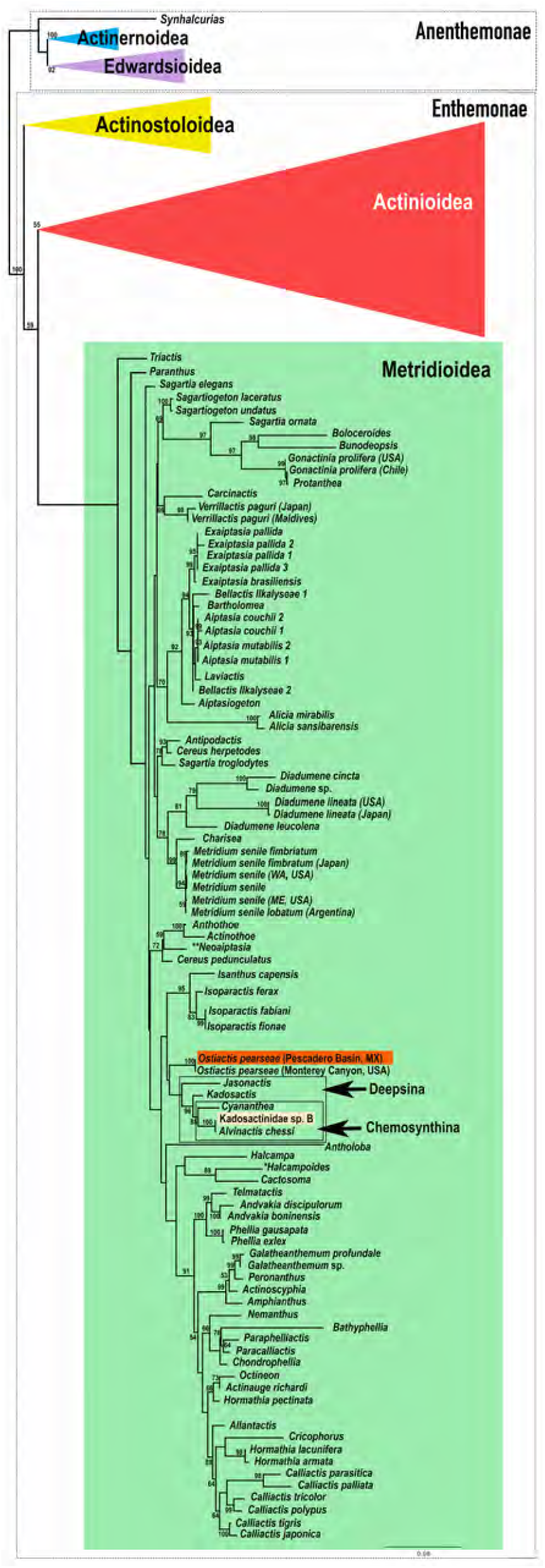
Phylogenetic placement of *Ostiactis pearseae*. Phylogenetic reconstruction resulting from maximum likelihood analysis using PhyML (RaxML results not shown, but congruent) of the concatenated dataset of three mitochondrial (12S rDNA, 16S rDNA, COIII) and a partial nuclear marker (18S rDNA). Doted boxes indicate actiniarian suborders; colored triangles and green box indicate actiniarian superfamilies; empty boxes and arrows indicate relevant actiniarian clades. Position of *Ostiactis pearseae* specimens from Pescadero Basin vent communities is highlighted in the orange box; the position of an additional sea anemone (unidentified morphospecies Kadosactinidae ‘sp. B’) is indicated by the light orange box. Bootstrap resampling values indicated above branches; only support values > 50% are shown.

**Table 2.** Size ranges of the cnidae capsules of *Ostiactis pearseae* (Daly & Gusmão, 2007). N: total number of capsules measured. F: Frequency, +++ = very common, ++ = common, + = rather common, * = sporadic.

| Categories | Range length<br>× width (µm) | Avg ± SD | N | S | F | Range length<br>× width (µm)<sup>3</sup> |
| --- | --- | --- | --- | --- | --- | --- |
| COLUMN<sup>1</sup> |  |  |  |  |  |  |
| Basitrichs | 17.2-26.0 x 2.6-4.4 | 21.7±2.3 x 3.4±0.3 | 122 | 6/6 | ++ | 13.1–22.5 x 1.9–3.0 |
| Holotrichs | 16.9-24.7 x 4.8-8.6 | 21.5±2.0 x 6.4±0.8 | 67 | 4/6 | */++ | 16.6–27.0 x 3.5–5.5 |
| *p*-mastigophores B1 | ---- | ----- |  | 0/6 |  | 17.8–33.9 x 2.6–5.3 |
| TENTACLES |  |  |  |  |  |  |
| Robust spirocysts | 15.9-41.9 x 4.2-8.8 | 26.0±6.1 x 5.7±1.0 | 85 | 5/5 | ++ | 16.3–35.5 x 2.1–5.8 |
| Basitrichs | 15.2-33.2 x 2.3-4.3 | 23.3±4.0 x 3.5±0.5 | 113 | 5/5 | +++ | 16.2–29.8 x 2.2–4.6 |
| Holotrichs | 15.6-28.5 x 4.0-9.3 | 23.3±2.7 x 7.1±1.0 | 79 | 5/5 | + | 23.1–34.4 x 2.9–6.9 |
| ACTINOPHARYNX |  |  |  |  |  |  |
| Basitrichs 1 | 16.4-20.2 x 2.6-3.2 | 18.1±1.7 x 2.9±0.2 | 5 | 2/3 | * | 16.8–26.3 x 2.0–3.3 |
| Basitrichs 2 | 23.7-34.0 x 3.3-4.6 | 27.9±2.8 x 4.0±0.4 | 20 | 2/3 | ++ | 25.6–44.5 x 3.0–4.3 |
| *p*-mastigophores A<sup>2</sup> | 23.3-38.1 x 4.7-6.9 | 30.7±3.5 x 5.7±0.5 | 53 | 3/3 | ++ | ----- |
| *p*-mastigophores B1<sup>2</sup> | 14.5-19.5 x 4.0-5.2 | 17.8±1.7 x 4.5±0.4 | 13 | 3/3 | ++ | 22.0–37.0 x 3.9–6.2 |
| FILAMENTS |  |  |  |  |  |  |
| Basitrichs | 15.3-23.3 x 2.5-3.5 | 18.5±1.6 x 3.0±0.2 | 50 | 4/4 | */+ | 14.1–22.5 x 1.8–2.9 |
| *p*-mastigophores A<sup>2</sup> | 26.5-37.8 x 4.8-6.7 | 32.8±2.2 x 5.7±0.5 | 49 | 4/4 | + |  |
| *p*-mastigophores B1<sup>2</sup> | 13.4-21.3 x 3.6-6.3 | 17.4±1.7 x 4.6±0.6 | 59 | 4/4 | +++ | 17.5–37.6 x 3.4–6.0 |

<sup>1</sup>Two specimens with papillae in distal column (see Fig. 2); papillae with only basitrichs of similar sizes than those in the rest of the column (i.e. 17.0-21.6 x 2.7-3.7 µm).
<sup>2</sup>Categories pooled together as microbasic *p*-mastigophores in Daly & Gusmão (2007).
<sup>3</sup>Data from Daly & Gusmão (2007).

Diagnosis: (amended after Daly and Gusmão, 2007 and Rodríguez et al. 2012, modifications in italics). Ostiactinidae with basilar muscles and mesogleal marginal sphincter. Body with well-developed base. Column not clearly divisible into scapus and scapulus; scapus without cuticle, *maybe* with scattered demarcated suckers distally; column without cinclides *or with a distal row of round papillae with inconspicuous cinclides.* Tentacles regularly arranged, not thickened on the aboral side. Six pairs of perfect and fertile mesenteries, hexamerously arranged, not divisible into macro- and micro-cnemes. *Same number of mesenteries proximally and distally.* Retractor muscles weak but *restricted*. No acontia. *Some populations with chemosynthetic bacteria in tentacles.* Cnidom: Robust spirocysts, basitrichs, holotrichs, and *p-mastigophores A and B1. Ostiactis pearseae* had been previously collected only in deep-sea waters (2800 m depth) of the Eastern Pacific, at a whalefall habitat in Monterey Bay (Daly & Gusmão 2007). Newly collected specimens are from deep-sea waters (3655-3692 m depth) associated with Southern Pescadero Basin hydrothermal vents (Diane’s vent) in the Gulf of California (Pacific Ocean).

Intrapopulation variability in morphology was observed in the Pescadero Basin *Ostiactis pearseae* specimens (Fig. 2). Differences were observed mainly in the abundance and categories of cnidae among specimens (particularly holotrichs in the column), but also in the presence of a distal row of papillae (with only basitrichs) associated with inconspicuous cinclides in two specimens (SO197-S2 and SO200-R2; Fig. 2). Because of the small size of the papillae, the relatively small sizes and state of contraction and preservation condition of the specimens, it is not definitive that this row of distal papillae is only present in these two individuals. Nevertheless, the rest of the morphological and molecular characters, as well as the cnidae data, from the specimens with distal papillae agree with those of the other specimens studied, suggesting that differences should be treated as intrapopulation variation. Morphologically, specimens of *O. pearseae* from the Pescadero Basin possess ~70 tentacles, compared to specimens of similar sizes from the type locality, described as having ~100 tentacles (Daly & Gusmão 2007), with some having poorly demarked suckers in the column (which were not observed in Pescadero Basin specimens). Additionally, the first and second cycles of mesenteries are fertile in the type specimens, with males observed brooding larvae internally in the tentacles (Daly & Gusmão 2007). Although the fertility of the first cycle could not be confirmed for the Pescadero Basin specimens, the second and third cycles were confirmed to be fertile, but no brooding individuals were identified. The previous implementation of a different cnidae terminology suggests conspicuous differences in cnidae types and sizes between specimens at the Monterey Canyon whalefall and Pescadero Basin (Table 2), but a new more precise combined terminology used here allows for distinction within *p*-mastigophore capsules (i.e. *p*-mastigophores A and *p*-mastigophores B1). The types and size ranges of the original description and the newly collected specimens of *O. pearseae* mostly agree (with only slight variability in some size ranges), with the only distinct difference being the presence of *p*-mastigophores B1 capsules in the tentacles of the whalefall specimens, and not the Pescadero Basin specimens (Table 2).

All molecular phylogenetic analyses, based on the concatenated mitochondrial 12S rDNA, 16S rDNA, COIII genes and partial nuclear 18S rDNA gene were congruent and revealed a well-supported clade comprised of specimens of *Ostiactis pearseae* from Pescadero Basin and the type locality of Monterey Canyon (Fig. 3). DNA sequences from the Pescadero Basin specimens were identical to the Monterey Canyon population for 3 of the genes analyzed, and only differed from the type locality by 1-bp for the 16S rDNA gene. *Ostiactis* was recovered within Metridioidea, as sister to a weakly supported clade formed by deep-sea Actiniaria and those associated with chemosynthetic environments (e.g. clades Deepsina + Chemosynthina, as part of the family Kadosactinidae, sensu Rodríguez et al. 2012), a relationship consistent over different studies (e.g. Rodríguez et al. 2012, 2014; Grajales and Rodríguez 2016; Gusmão et al. 2019). Most representatives from these two clades are characterized by the loss of acontia (filament-like structures packed with nematocysts), the presence of which is a mayor synapomorphy for Metridioidea. The other actiniarian morphotype included here, identified as Kadosactinidae ‘sp.B’, was recovered (only mitochondrial sequence data available for this specimen) sister to *Alvinactis chessi*, a sea anemone inhabiting hydrothermal vents in the southwestern Pacific (Fig. 3; Zelnio et al. 2009).

### *Isotope signatures of* Ostiactis pearseae *from the Pescadero Basin*

Tissue stable δ^13^C, δ^15^N and δ^34^S isotope values were significantly different for *Ostiactis pearseae* than other anthozoans at the Pescadero Basin vent fields (ex. zoanthids; Fig. 4A). For example, *O. pearseae* had δ^13^C tissue values of −29.1 ± 4.6‰ (n = 9), while the others measured −20.6 ± 2.7‰ (n = 10; ± 1 SD; ANOVA p = 0.0001; Fig. 4A). Similarly, *O. pearseae* had much more negative δ^15^N tissue values of 1.6 ± 1.7‰, whereas the others measured 10.6 ± 6.3‰, a difference of ~9‰ (± 1 SD; ANOVA p = 0.0006; Fig. 4A). Low δ^15^N values were also observed in nearly all individuals of the Kadosactinidae ‘sp.B’ (~0.5-2.8; Fig. 4A) collected at the same locality. By comparison, *O. pearseae* was significantly more negative in both tissue δ^13^C and δ^15^N than four methane seep-associated octocoral species recently reported on by Vohsen et al. 2020 (ANOVA p = 0.0013 and p < 0.00001, respectively, n = 21; Fig. 4A). Finally, *O. pearseae* had significantly lower δ^34^S tissue values of −1.1 ± 6.4‰ (n = 7), compared to other Pescadero Basin sea anemones (8.9 ± 5.2‰; n = 6; ANOVA p = 0.008), with comparable total tissue sulfur ~ 0.9-1.9% by weight (Fig. 4B).

**Figure 4:**
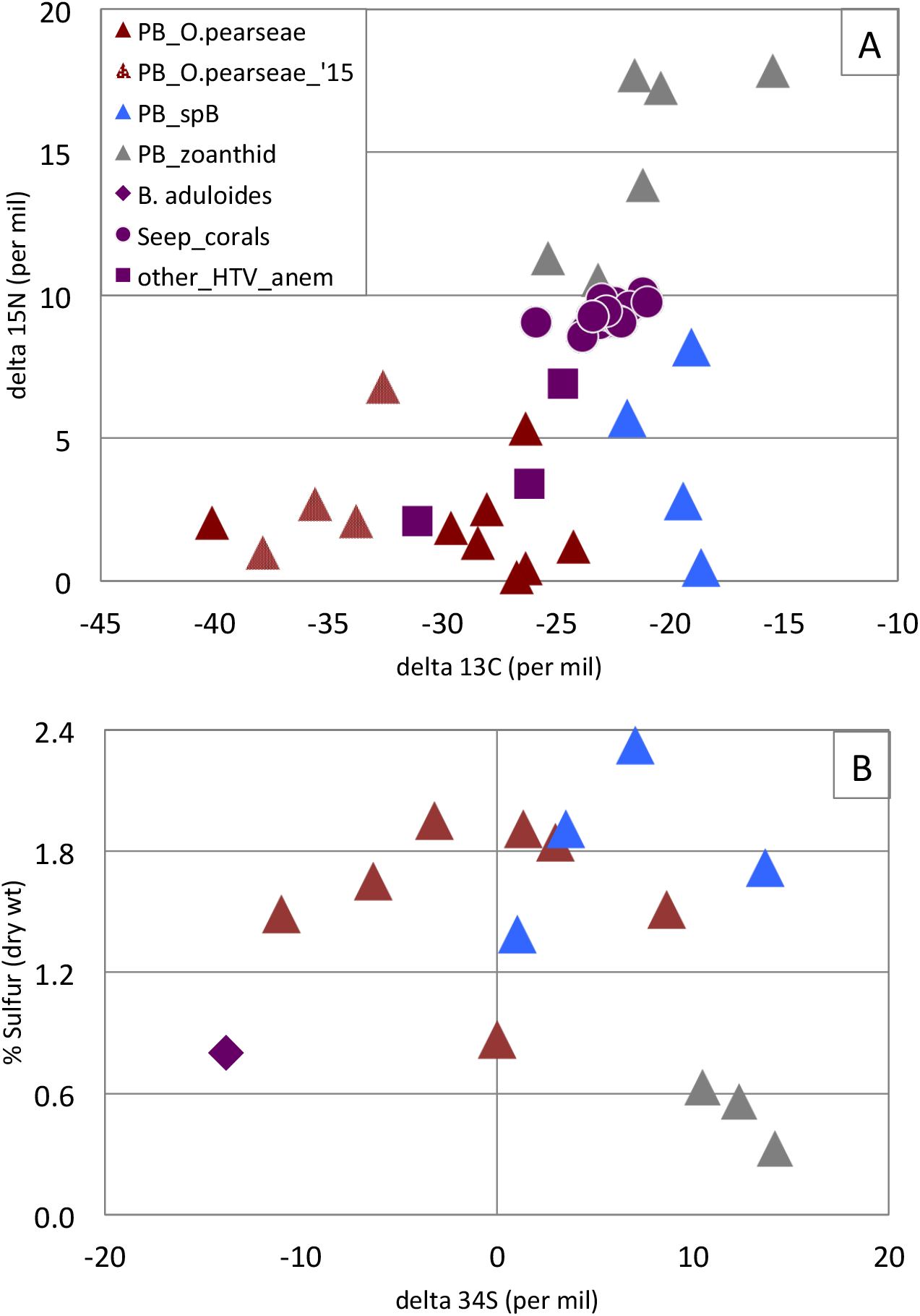
^13^Carbon, ^15^Nitrogen, ^34^Sulfur isotope signatures for *Ostiactis pearseae* and comparison actiniarians. **A.** δ^13^C and δ^15^N values for the tentacles of *Ostiactis pearseae* from the Pescadero Basin vents, compared to neighboring anthozoans, including unidentified zoanthids (‘zoan’) and sea anemone (Kadosactinidae ‘sp. B’). Data for Pescadero Basin actiniarians collected in 2015 (from Goffredi et al. 2017; red checkered triangles) as well as seep-associated corals from the Gulf of Mexico (Vohsen et al. 2020; purple circles) and unidentified anemones from Gorda Ridge hydrothermal vents (Van Dover & Fry 1994; purple squares) are also included. Data from Vohsen et al. 2020 was extracted from their Figure 7 using an online Web plot digitizer (http://arohatgi.info/WebPlotDigitizer/). **B.** δ^34^S and tissue sulfide content (%, dry weight) values for the tentacles of *O. pearseae* from the Pescadero Basin vents, compared to neighboring Anthozoa, including an unidentified zoanthid (‘zoan’) and sea anemone (Kadosactinidae ‘sp. B’). *Bathymodiolus aduloides* from muddy sediments off of Kakaijima Island taken from Yamanaka et al. 2000.

### *Bacterial community analysis of* Ostiactis pearseae *from the Pescadero Basin*

In *Ostiactis pearseae*, the tentacles are smooth, tapering and relatively long when extended, with features similar to most other anemones, including cnidocytes (or cnidoblasts), cnidocysts and glandular cells, all common cellular components sea anemones tentacles (Fautin & Mariscal 1991). Daly & Gusmão (2007) did not detect the presence of bacteria in the type specimens of *O. pearseae* from Monterey Canyon, however, the unusual isotope signatures of the Pescadero Basin specimens (Goffredi et al. 2017; Salcedo et al. 2019) prompted a more careful examination of this possibility. Indeed, bacterial community analysis via 16S rRNA Illumina barcoding revealed a dominance of the gammaproteobacteria SUP05 clade (64-96% of the bacterial community), comprising 6 putative sulfide-oxidizing bacterial OTUs (= phylotypes, clustered at 99% similarity) associated with the tentacles of Pescadero Basin *O. pearseae* (n = 8 specimens; Fig. 5, Table S1). This was in contrast to the sulfide-oxidizing gammaprotebacteria recovered from the nearby obligate vent tubeworms *Riftia pachyptila* and *Oasisia* aff. *alvinae* (100% identical to Candidatus *Endoriftia persephone;* Fig. 5A; Robidart et al. 2008). The SUP05 clade was not detected in association with 6 other individual sea anemones (determined by morphotype or molecular sequencing) and the specific *Ostiactis*-associated SUP05 phylotypes were not detected in surrounding water samples (n = 3; Fig. 5). Three SUP05 OTUs comprised 5-7% of the bacterial community in the surrounding water column (Fig. 5A), but were distinct, based on the 250-bp 16S rRNA Illumina barcode sequences (Fig. S1 inset).

**Figure 5:**
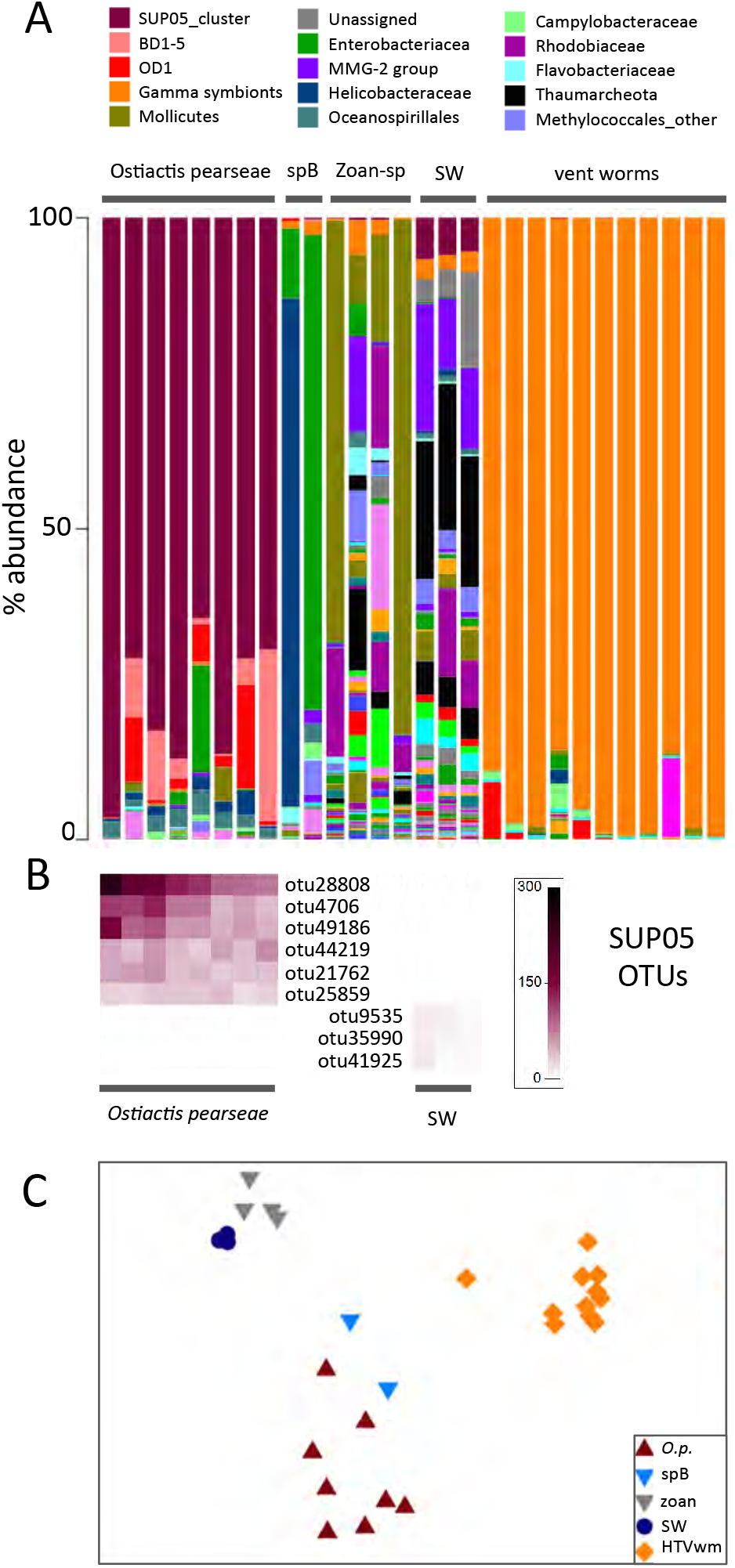
Relative abundance of 16S rRNA bacterial phylotypes, recovered from *Ostiactis pearseae* and comparison samples. **A.** Relative abundance of bacterial families from *Ostiactis pearseae* from the Pescadero Basin vents, compared to neighboring Anthozoa, including unidentified zoanthids (‘zoan’) and Kadosactinidae ‘sp. B’, nearby obligate vent tubeworms *Riftia pachyptila* and *Oasisia* aff. *alvinae*, and seawater. Each color on the graph represents a distinct family-level phylotype or lowest level available. The top 15 dominant family phylotypes are indicated in the key. For all others, see DataFile S1, including raw and processed data, as well as representative sequences for all dominant hits. **B.** Six distinct SUP05 OTUs (99% 16S rRNA sequence similarity) recovered from *O. pearseae*, compared to the surrounding seawater. The heatmap scale reflects the number of reads per sample. Phylogenetic relationships between the SUP05 OTUs are shown in Figure S1. See also DataFile S1, for representative sequences of each OTU. **C.** Non-metric multidimensional scaling (NMDS) ordination of microbial communities associated with *O. pearseae*, versus the other neighboring species and overlying seawater. Each point represents all 16S rRNA sequences recovered from a single specimen or sample. ANOSIM p < 0.022, suggesting a significant difference between *O. pearseae* and any other sample set (ex. other sea anemones, water samples; R = 0.88-1.00). HTV = hydrothermal vent. spB = another undescribed sea anemone (Kadosactinidae ‘sp. B’) from the Pescadero Basin. zoan = unidentified zoanthids from the Pescadero Basin.

NMDS ordination revealed the total bacterial community of *Ostiactis pearseae* to be strongly differentiated from those associated with zoanthid specimens (Analysis of Similarity (ANOSIM) R = 0.99, p = 0.002), the other unidentified sea anemone Kadosactinidae ‘spB’ (ANOSIM R = 0.87, p = 0.022), the water column samples (ANOSIM R = 1.00, p = 0.006), and the neighboring obligate vent tubeworms *Riftia pachyptila* and *Oasisia* aff. *alvinae* (ANOSIM R = 1.00, p = 0.001; Fig. 5C). Bacterial community analysis revealed limited diversity within the tentacles of *O. pearseae* from the 5 different Pescadero Basin vent sites (Fig. 5A; Table S1). Other bacteria uniquely recovered from *O. pearseae* tentacles included the BD1-5 group (a.k.a. Gracilibacteria; present in 5/8 *O. pearseae* specimens at 3-27%) and the OD1 group (a.k.a. Parcubacteria; present in 5/8 *O. pearseae* specimens at 1-16%; Fig. 5A). Additional bacterial groups present in non-*Ostiactis* sea anemones included Enterobacteriacea and Mollicutes (the latter 89% similar to one recovered from an ascidian; Fig. 5A; EF137402; Tait et al. 2007). Microbial groups that were more common in all 3 water samples included the Methylococcales marine group 2 (MMG-2), Rhodobiacea, and Thaumarcheota (Fig. 5A).

To further characterize the SUP05 in association with *Ostiactis pearseae*, a longer region of the 16S rRNA gene was amplified via direct PCR and sequenced. A 1334-bp long 16S rRNA sequence, only 1-bp different from barcode OTU21762, was 96.5% similar to a free-living bacterium from a mud volcano in the Eastern Mediterranean Sea (AY592908; Heijs et al. 2005) and 95% similar to the thiotrophic symbiont of *Bathymodiolus* aff. *brevior* from Central Indian Ridge vents (DQ077891; McKiness and Cavanaugh 2005; Fig. S1). There are no *Bathymodiolus* mussels at the Pescadero Basin vents, and the SUP05-related sequences recovered from *O. pearseae* are distinct from those recovered from *Bathymodiolus* from Costa Rica seeps, some of the closest known mussel populations (only ~96% similar for 250bp barcode sequence; Fig. S1 inset; Levin et al. 2012; McCowin et al. in press).

Several genes were additionally amplified and directly sequenced in order to inform the possible metabolic capabilities of the SUP05-related bacteria in *O. pearseae.* The *napA* gene, encoding a catalytic subunit of the periplasmic nitrate reductase alpha subunit (E.C. 1.7.99.4; Flanagan et al. 1999), amplified from *O. pearseae* tentacles, was most closely related (82-85% similarity based on amino acid translation; 77% based on nucleotides), to the *napA* gene from known SUP05 bacteria, including those from an estuary (ACX30474; Walsh et al. 2009) and the endosymbiont from *Bathymodiolus* mussel gill tissues (SMN16186). The *aprA* gene, encoding the adenosine phosphosulfate (or APS) reductase alpha subunit (E.C.1.8.99.2), recovered from *O. pearseae*, was most closely related to the *aprA* gene from the bacterial symbiont of a nematode from (96% similarity based on amino acid translation, ACF93728) and the endosymbiont of *Bathymodiolus septemdierum* (81% similarity based on nucleotide sequence, AP013042; Fujiwara et al. 2000).

Abundant SUP05 bacteria were observed embedded within the tentacle epidermis of *Ostiactis pearseae* (Fig. 6). Fluorescent *in situ* signal amplification via hybridization chain reaction-FISH (HCR-FISH) was necessary to overcome the very highly autofluorescent cnidae produced by the epidermis (Fig. 6G). HCR-FISH and TEM microscopy revealed intracellular cocci-shaped cells (~0.5 μm diameter), positioned just above or immediately adjacent to cnidae capsules and nuclei (Fig. 6G,I-J). These cells were definitively identified as members of the SUP05 group, given the consistent overlap between cells hybridized using a general bacterial probe set (Eub338-I-III) and a probe designed specifically to target the *O. pearseae* SUP05 (Fig. S2). Although poor tissue fixation somewhat compromised high-resolution electron microscopy (ex. many host cells were visibly ruptured with jumbled mitochondria), TEM provided additional evidence of symbiont integration within *O. pearseae.* Bacteria within the tentacles appeared concentrated in the periphery of cells within the mono-layered epidermis (Fig. 6I). Additionally, glands with large electron dense vesicles were observed occasionally between the very elongated bacteria-containing cells (Fig. 6G) and a layer of mucous was observed overlying the epidermis in some instances. Bacteria on the tentacle surface occasionally appeared to be in clathrin-coated pits in various stages of endocytosis (Fig. 6L). For both microscopy methods, bacteria were not observed in either the gastrodermis or mesoglea of *O. pearseae* (Fig. 6G).

**Figure 6.**
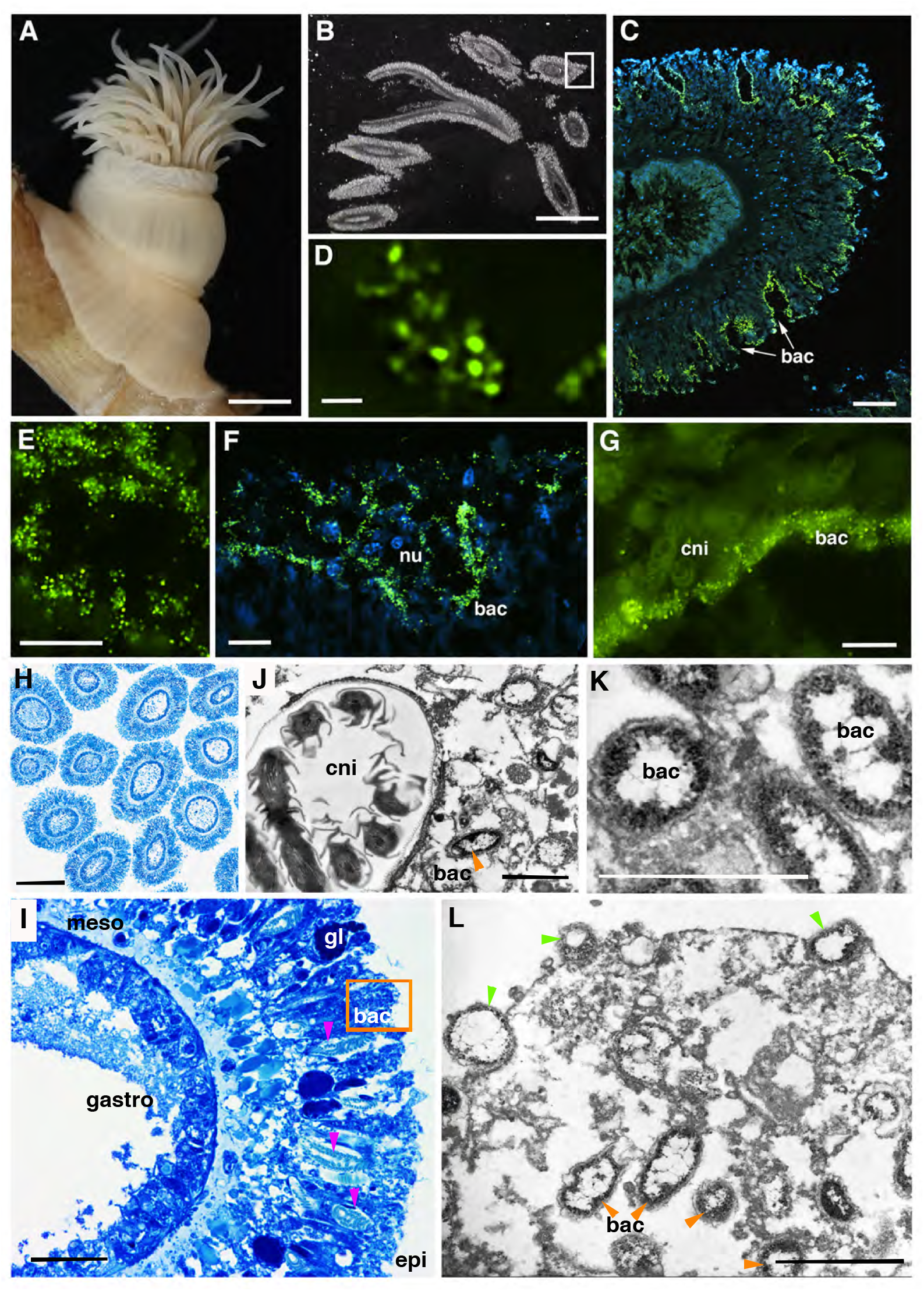
Microscopy of the tentacles of *Ostiactis pearseae*. **A**. Whole specimen image - SO194-S2, SIO-BIC Co3067. **B.** light microscopy of 3-μm sections embedded in Steedman’s resin, and **C-G**. fluorescent *in situ* signal amplification via hybridization chain reaction-FISH (HCR-FISH) microscopy of *Ostiactis pearseae* tentacles. An unlabeled probe (Anem_SUP05), with a specific sequence initiator tag was designed to be an exact match to the putative thiotrophic symbiont (related to the SUP05 clade). This probe was then amplified via HCR-FISH using DNA hairpins labelled with Alexa488, shown in green. DAPI-stained nuclei of host cells are shown in blue. **F-G.** Bacteria can be seen within the epidermis, in and amongst nuclei, positioned just above or immediately adjacent to cnidocysts. **H-I**. Light microscopy of *O. pearseae* tentacles**. J-L**. Transmission electron (TEM) microscopy of *O. pearseae* tentacles. **I.** Bacteria are concentrated in the periphery of elongated epidermal cells (designated by the orange box, which corresponds to the area of TEM imagery), and positioned near cnidae, shown in pink arrowheads. No bacteria were observed in the mesogloea or gastrodermis. **J.** Close-up of bacteria near a cnidocyst capsule, with enclosed tubule. **K.** Close-up showing clear membranes surrounding the bacterial cells, designated by orange arrowheads. **L.** Arrowheads (in green) point to bacteria possibly being endocytized via clathrin-coated pits, as well as nearby clusters of bacterial cells within the elongated epidermal cells of *O. pearseae.* nu, nucleus. bac, bacteria. cni, cnidae. meso, mesoglea. gastro, gastrodermis. epi, epidermis. Scale bars are 5 mm (A), 2 mm (B), 50 μm (C), 1 μm (D), 10 μm (E-G), 250 μm (H), 25 μm (I), 1 μm (J-L).

## Discussion

A conspicuous actiniarian species, identified as *Ostiactis pearseae* (Daly & Gusmão, 2007), was dominant at two neighboring hydrothermal vent fields in the Pescadero Basin, Gulf of California. Unlike most vent anemones, which are almost always observed in the vent periphery, this species was found very near to vigorous venting fluids on and among the obligate vent tubeworms *Oasisia* aff*. alvinae* and *Riftia pachyptila* (Fig. 1), known to rely exclusively on sulfide-based chemosynthesis for energy (Fisher et al. 1989; Van Dover & Fry 1989). *Ostiactis pearseae*, formerly named *Anthosactis pearseae* (see Rodríguez et al. 2012), had been originally described as the first and only endemic Actiniaria from a whalefall community (Daly & Gusmão 2007), however, this discovery at hydrothermal vents makes them one of the only sea anemones described from multiple chemosynthetic environments (Zelnio et al. 2009; Rodríguez et al. 2012). The assumption, until now, was that most sea anemones at hydrothermal vents and methane seeps acquire nutrients via suspension feeding. Daly & Gusmão (2007) previously found no evidence that *Ostiactis pearseae* harbored chemosynthetic bacteria and accepted that they fed upon dissolved and particulate organic matter and plankton. However, the significantly negative δ^13^C and δ^15^N tissue isotopic values of *O. pearseae* (at the time labeled as an unidentified species in Goffredi et al. 2017 and Salcedo et al. 2019), suggested an entirely different strategy dependent upon bacteria chemosynthesis.

Indeed, distinct bacterial phylotypes related to the SUP05-group were associated with *Ostiactis pearseae*, compared to other nearby sea anemones and water column bacterial communities. This association was pervasive and dominant, in that SUP05 bacteria were found in all 8 *O. pearseae* specimens analyzed, comprising up to 96% of the recovered microbial 16S rRNA genes. Sulfur-oxidizing bacteria within the SUP05 clade, named after discovery in the Suiyo seamount plume (Sunamura et al. 2004), have been found worldwide in marine oxygen-deficient marine environments, deep-sea hydrothermal systems, and productive upwelling regions (Labrenz et al. 2007; Ulloa et al. 2012; Glaubitz et al. 2013). They exist both as free-living cells (Walsh et al. 2009) and in association with animal hosts (ex. *Bathymodiolus* mussels and some sponges; Petersen et al. 2012), where they participate centrally in the provisioning of fixed carbon to the animal.

Intracellular SUP05 were observed exclusively in the epidermis of *Ostiactis pearseae*, which was unexpected given that most Cnidaria house symbionts, mainly photosynthetic, in the gastrodermis (McAuley 1985; Marlow & Martindale 2007; Mellas et al. 2014). Epidermal bacteriocyte-like structures containing *Vibrio* have been observed in *Exaiptasia pallida* (Palincsar et al. 1989) and bacterial ‘aggregates’ containing *Endozoicimonas* have been observed in epidermal ‘caverns’ in the sea anemone *Metridium senile* (Schuett et al. 2007). In both cases, however, the epidermal bacteria are pathogens commonly associated with animals (Preheim et al. 2011; Neave et al. 2016). Vohsen et al. (2020) reported SUP05, and other bacteria, associated with whole octocoral specimens, including mucous, however the specific location of these bacteria was not determined. Interestingly, bacteria on the tentacle surface of *O. pearseae* appeared to be in clathrin-coated pits in various stages of endocytosis. Further examination of this receptor-mediated process is necessary to establish whether bacteria are actively transported inside of host cells and if so, what influences the recognition and selectivity of this process.

Hosting sulfide-oxidizing SUP05 in the outer epidermis may allow *Ostiactis pearseae* to avoid sulfide toxicity or the costly evolution of unique biochemistry to take up and transport sulfide (Goffredi et al. 1997). Additionally, the epidermis in Anthozoa can function in nutrition (Fautin & Mariscal 1991), even more so than the gastrodermis, through direct uptake of dissolved organic compounds (Schlichter 1975, 1980), thus the positioning of nutritional bacteria in the epidermis may increase effective exchange of small molecules. Like other SUP05 cells, those associated with the tentacles of *O. pearseae* were small (~500 nm in diameter; Shah et al. 2019). Presumably these symbionts require both oxygen and sulfide near simultaneously, for example *“Candidatus* Thioglobus autotrophicus”, a member of the SUP05 group, has an aerobic phenotype, and uses sulfide while respiring oxygen (Marshall and Morris 2013; Shah et al. 2019). In this regard, it would be reasonable to house bacteria as close to the tissue surface as possible in order to accommodate gas exchange and meet symbiont metabolic demands.

The assumption that the SUP05 group may perform a nutritional role for the Pescadero Basin *Ostiactis pearseae* is evidenced by the comparatively light tissue δ^13^C values (average −29.1‰). The contribution of chemosynthesis-derived carbon to *O. pearseae* biomass appears to exceed that reported for deep-sea anthozoan species from the Gulf of Mexico (ex. *Swiftia* and *Acanthogorgia*; Vohsen et al. 2020). The facultative nature of the SUP05-Anthozoa symbioses proposed by Vohsen et al. 2020 is also suggested for *O. pearseae* given the large range in negative δ^13^C values observed. Like all sea anemones (even those with photosynthetic symbionts), *O. pearseae* retains an arsenal of nematocysts by which to capture prey, thus the SUP05 symbionts likely provide only a portion of their diet.

The SUP05 clade is not only involved in mediating dark carbon fixation, but also the cycling of nitrogen, whether by denitrification, as has been shown in free-living SUP05 populations (Walsh et al. 2009; Glaubitz et al. 2013) or assimilatory nitrate reduction, as in the case of symbiotic SUP05 (Ikuta et al. 2016; Vohsen et al. 2020). Significant contribution to tissue nitrogen by microbial nitrate utilization may be possible for the SUP05 symbionts given the considerably low δ^15^N values in *O. pearseae* (average 1.6‰) and the successful amplification of the SUP05-related periplasmic nitrate reductase alpha subunit (*napA*) gene. The actual abundance of SUP05 symbionts per individual anemone is not known, nor is the regulation of carbon or nitrogen nutrient exchange, and thus the overall nutritional influence of the SUP05 bacteria is not yet quantifiable.

Finally, *Ostiactis pearseae* tissue δ^34^S values (~-1‰) represented a large offset from typical marine biomass (16-21‰; Kaplan et al. 1963), where biogenic sulfur is sourced from seawater sulfate with minimal isotopic fractionation (21‰; Paris et al. 2013). The average δ^34^S observed in *O. pearseae* tissues is consistent with typical hydrothermal vent fauna (−5 to +5‰; Fry et al. 1983), which are known to incorporate a local source of sulfur (e.g. volcanic, thermally-altered sulfur at ~0‰; Sakai et al. 1984; Canfield 2004) via internal symbioses or direct consumption of sulfide-oxidizing bacteria. However, several individuals of *O. pearseae* revealed even lower δ^34^S values (down to −11‰), which would likely require the additional incorporation of substantial sulfide produced via microbial sulfate reduction, which is expected to have a δ^34^S signature of −20‰ or lighter (Chambers & Trudinger 1979; Morse et al. 1987). The incorporation of sulfide sourced from dissimilatory sulfate reduction, rather than hydrothermal sulfide, has been similarly proposed for SUP05-hosting *Bathymodiolus* mussels from both Kakaijima Island and the Kaikata Caldera, which had tissue δ^34^S values of −12‰ and −25‰, respectively, with a comparable tissue sulfur content of ~0.8% (n = only 1 specimen each; Kim et al. 1989; Yamanaka et al. 2000). The wide range of δ^34^S values for *O. pearseae* tissues (~ −11 to 9‰), compared to other thiosymbiont-hosting animal species, could be due to a combination of traditional feeding by the host, variable sulfide oxidation by the SUP05 symbionts (e.g. utilization of H_2_S, HS^-^, or other reduced S species, including endogenous elemental sulfur), or variation in the sulfur sourced from the petroleum-rich sediments of the Pescadero Basin.

Many uncultivated candidate bacterial phyla have been discovered in recent years within a variety of environments (Rinke et al. 2013; Kantor et al. 2013; Harris et al. 2014). They usually have small genomes (<1 Mb) with dramatically reduced biosynthetic capabilities, and yet exist globally in both marine and terrestrial habitats (Wrighton et al. 2012). Several of these candidate phyla, known as the OD1 and BD1-5 groups (also referred to as Parcubacteria and Gracilibacteria, respectively), comprised up to 16-27% of the

*Ostiactis pearseae* bacterial community, and are known to play an important role in sulfur cycling (Wrighton et al. 2012). Nelson and Stegan (2015) proposed an ectosymbiotic or parasitic lifestyle for the OD1, given their inability to synthesize vitamins, amino acids, nucleotides, and fatty acids. Additionally, while most candidate phyla are found in anoxic habitats, some OD1 genomes contained genes suggestive of O2 use as a terminal electron acceptor (Brown et al. 2015; Nelson and Stegan 2015). Although not previously associated with the sulfide-oxidizing SUP05 group, or any specific proteobacterial group, the role of OD1 in sulfur reduction (Wrighton et al. 2012) and their diverse repertoire for attachment and adhesion (Nelson and Stegan 2015) forecasts a possible direct association with either the SUP05 bacteria or *O. pearseae* mucous, for example.

## Conclusion

Despite 40+ years of appreciation for chemosynthetic symbioses and the continued search for their occurrence in the most well-known habitats, Cnidaria have not been among the animals known to associate with chemoautotrophic bacteria. Here, we identify a hydrothermal vent sea anemone, *Ostiactis pearseae*, at 3700 m depth in the Pescadero Basin, Gulf of California, that appears to be nutritionally supported by internal chemoautotrophic bacteria. This species, one of only 2 dominant sessile animals observed on the vent chimneys, has an unusual life position, often located in and amongst vent-obligate siboglinid tubeworms, very near to actively venting fluids. *Ostiactis pearseae* houses putative sulfide-oxidizing SUP05 bacteria in its epidermis, with which it appears to have established a facultative nutritional symbiosis, based on a broad range of carbon, nitrogen, and sulfur isotopes. Facultative nutritional symbioses are often more difficult to recognize, compared to obligate alliances, but they are surely more common in nature (Goffredi et al. 2020), particularly in Cnidaria which experience symbiont gain and loss readily (ex. Jones et al. 2008; Larson et al. 2014; Vohsen et al. 2019). So, too, is the difficulty in uncovering nested symbioses, often involving microbe-microbe synergies. In this study, an unusual abundance of two candidate phyla, Parcubacteria and Gracilibacteria (a.k.a. OD1 and BD1-5, respectively) within *O. pearseae* tentacles, hints at the roles they may play in the cycling of nutrients within and on animal hosts. Cnidaria symbioses are considered foundational for coral reefs, and perhaps they also play an important role at hydrothermal vents. It would be worth investigating additional Anthozoa species observed to inhabit venting fluids at other sites worldwide (Doumenc & Van-Praët 1988; Sanamyan & Sanamyan 2007; Rogers et al. 2012), to see whether they have also forged nutritional relationships with chemosynthetic bacteria, such as the versatile SUP05 group.

## Methods

### Specimen Collections

All specimens and water samples were collected from active vent sites within the Pescadero Basin, Gulf of California (~ 3700 m depth), using the ROV *SuBastian* during the R/V *Falkor* expedition FK103118 (October-November 2018), specifically from six sites at two vent fields within ~2 km of each other; the previously described Auka vent field (refs) and a newly discovered JaichMaa ‘ja’ag vent field (Fig. 1; Table 1). Sea anemones were collected by ROV manipulator or suction sampler (Supplemental video, currently available at https://doi.org/10.5061/dryad.mkkwh70wt) and preserved shipboard as described below in each analysis section. Targeted water samples (2 L) were collected via Niskin bottle mounted on ROV *SuBastian.*

### *Material examined for redescription of* Ostiactis pearseae

SIO-BIC Co3060 [GC18-0004] (S0193-R2): Specimens: 2; Details: Fixative 4% paraformaldehyde; Preservative: 50% EtOH; Matterhorn, Auka Vent Field, Pescadero Basin, Mexico (23.95404°N, 108.86296°W); 3655 m; 14-Nov-2018. SIO-BIC Co3061 [GC18-0005] (S0193-A2): Specimens: 1; 10% formalin, preserved 50% EtOH; Matterhorn to Diane’s Vent, Auka Vent Field, Pescadero Basin, Mexico (23.95472°N, −108.86233°W); 3655 m; 14-Nov-2018. SIO-BIC Co3067 [GC18-0028] (S0194-S2): Specimens: 2; fixed: 10% formalin; 50% EtOH; Z Mound, Auka Vent Field, Pescadero Basin, Mexico (23.95666°N, −108.86171°W); 3670 m; 15-Nov-2018. Material studied has been deposited in the Benthic Invertebrate Collection of Scripps Institution of Oceanography (University of California San Diego) and the Invertebrate Division collection of the American Museum of Natural History (AMNH) in New York.

Additional specimens examined in this study include Kadosactinidae ‘sp.B’ (SIO-BIC Co3065 [GC18-0012] (S0193-S4) and the unidentified zoanthid (SIO-BIC Co3066 [GC18-0025] (S0194-S1).

### Carbon, nitrogen, and sulfur isotope analysis

Tissue samples were dissected at sea, rinsed in milli-Q water, and frozen at −20°C until thawed, washed with milli-Q water, and dried for 48 h at 60°C. Carbon and nitrogen isotope determinations of anemone tissues were made via isotope ratio mass spectrometry. Samples (0.2-0.8 mg dry weight) were loaded in tin boats and analyzed for total organic carbon (TOC) and total nitrogen (TN) abundances and δ^13^C_org_ and δ^15^N using a Flash 2000 Elemental Analyzer (Thermo Fisher Scientific) interfaced to a Delta V Plus IRMS (Thermo Fisher Scientific) at Washington University, Missouri, USA. Samples were interspersed with several replicates of both in-house standards and international reference materials, including: IAEA-CH-6, IAEA-CH-3, IAEA-NO3, USGS-40, and USGS-41. TOC and TN abundances were quantified by integrating peak areas against those produced by in-house standards across a range of masses. The isotopic values are expressed in permil (‰) relative to international standards V-PDB (Vienna Pee Dee Belemnite) and Air for carbon and nitrogen, respectively. The long-term standard deviation is 0.2‰ for δ^13^C_org_ and 0.3‰ for δ^15^N. Sulfur isotope analyses were performed by combusting ~2-5 mg (dry weight) of tissue using a Costech ECS 4010 elemental analyser coupled to a Thermo Fisher Scientific Delta V Plus mass spectrometer. Sulfur isotope values are expressed in standard delta notation (δ^34^S) in permil (‰) as a deviation from the Vienna Canyon Diablo Troilite (VCDT) standard. The long-term standard deviation is 0.3‰ for δ^34^S is 0.3‰ based on in-house and international standards, including NBS-127 and IAEA-S1.

### DNA extraction

Specimens for molecular analysis (Table 1) were preserved immediately upon collection in ~90% ethanol and stored at 4°C. Total genomic DNA was extracted from tissues using the Qiagen DNeasy kit (Qiagen, Valencia, CA, USA) according to the manufacturer’s instructions. 2L water samples were filtered onto a 0.22 μm Sterivex-GP polyethersulfone filter (Millipore-Sigma, St. Louis, MO, USA) and frozen at −80°C until DNA analysis. DNA extraction from Sterivex PES filters was also performed using the Qiagen DNeasy kit, according to the manufacturer’s instructions, with the exception of the first step where 2 ml of ATL lysis buffer was added to the Sterivex filter, via luer lock and syringe, and rotated at 56°C for 12 hours. This solution was recovered from the filter, also via luer lock and syringe, and processed as usual.

### Molecular Analysis of the microbial community

A 1000-bp region of the gene coding for *napA* (periplasmic nitrate reductase) was amplified directly from *Ostiactis* tissues using the primers V16F (5’-GCNCCNTG-YMGNTTYTGYGG-3’) and V17R (5’-RTGYTGRTTRAANCCCATNGTCCA-3; Flanagan et al. 1999), while a 408 bp fragment of the *aprA* gene (subunit of particulate methane monooxygenase enzyme) was generated using primers, aps1F (5-TGGCAGATCATGATYMAYGG-3) and aps4R (5-GCGCCAACYGGRCCRTA-3, described in Blazejak et al. 2006). A 1465-bp fragment of the 16S rRNA gene was amplified using the primers 27F and 1492R. Annealing conditions of 50°C, 50°C and 54°C were used for *napA, aprA*, and 16SrRNA, respectively. Otherwise, all thermal protocols included the following steps: an initial 5 min denaturation at 94°C, then 1 min at 94°C, 1 min annealing step, and 1 min at 72°C, for 25 cycles, and a final 5 min extension at 72°C. Amplification products were sequenced directly using Sanger sequencing, via Laragen Inc., and submitted to GenBank (accession numbers XXXXXXX – TBD; currently available at https://doi.org/10.5061/dryad.mkkwh70wt). Close environmental and cultured relatives were chosen using top hits based on BLAST (www.ncbi.nlm.nih.gov).

The V4-V5 region of the 16S rRNA gene was amplified using bacterial primers with Illumina (San Diego, CA, USA) adapters on the 5’ end 515F (5’-TCGTCGGC-AGCGTCAGATGTGTATAAGAGACAGGTGCCAGCMGCCGCGGTAA-3’) and 806R (5’-GTCTCGTGGGCTCGGAGATGTGTATAAGAGACAGGGACTACHV-GGGTWTCTAAT-3’) (Caporaso et al. 2011). The PCR reaction mix was set up in duplicate for each sample with Q5 Hot Start High-Fidelity 2x Master Mix (New England Biolabs, Ipswich, MA, USA) and annealing conditions of 54°C for 25 cycles. Duplicate PCR samples were then pooled and 2.5 μL of each product was barcoded with Illumina NexteraXT index 2 Primers that include unique 8-bp barcodes (P5 5’-AATGATACGGCGACCACCGAG-ATCTACAC-XXXXXXXX-TCGTCGGCAGCGTC-3’ and P7 5’-CAAGCAGAA-GACGGCATACGAGAT-XXXXXXXX-GTCTCGTGGGCTCGG-3’). Secondary amplification with barcoded primers used conditions of 66°C annealing temperature and 10 cycles. Products were purified using Millipore-Sigma (St. Louis, MO, USA) MultiScreen Plate MSNU03010 with a vacuum manifold and quantified using Thermo-Fisher Scientific (Waltham, MA, USA) QuantIT PicoGreen dsDNA Assay Kit P11496 on the BioRad CFX96 Touch Real-Time PCR Detection System. Barcoded samples were combined in equimolar amounts into single tube and purified with Qiagen PCR Purification Kit 28104 before submission to Laragen (Culver City, CA, USA) for 2 x 250bp paired end analysis on the Illumina MiSeq platform with PhiX addition of 15-20%.

MiSeq 16S rRNA sequence data was processed in Quantitative Insights Into Microbial Ecology (v1.8.0). Raw sequence pairs were joined and quality-trimmed using the default parameters in QIIME. Sequences were clustered into *de novo* operational taxonomic units (OTUs) with 99% similarity using UCLUST open reference clustering protocol, and then, the most abundant sequence was chosen as representative for each *de novo* OTU. Taxonomic identification for each representative sequence was assigned using the Silva-119 database, clustered at 99% similarity. A threshold filter was used to remove any OTU that occurred below 0.01% in the combined samples dataset. Analyses are based on Bray-Curtis distances of fourth-root transformed data, which minimizes errors in the ordination due to PCR bias, while not sacrificing genuine differences between samples. Quantification and statistical analyses are described in the Results sections and figure legends. Comparisons were performed using ANOVA and statistical significance was declared at P < 0.05. Statistical analyses of beta diversity (e.g. ANOSIM) were performed with Primer E. The raw and processed Illumina 16S rRNA sequence data, as well as representative sequences, are available in DataFile S1 on the Dryad Digital Repository (https://doi.org/10.5061/dryad.mkkwh70wt).

### *Molecular Analysis of the anemone host* Ostiactis pearseae

Phylogenetic relationships were determined via sequencing of three mitochondrial markers, the 12S rRNA, 16S rRNA, and cytochrome oxidase III genes, and the partial nuclear 18S rRNA gene. An 862-bp product of the 12S rRNA gene was amplified via primers ANTMT12SF (5’-AGCCAC-ACTTTCACTGAAACAAGG-3’) and ANTMT12SR (5’-GTTCCCYYWCYCTYA-CYATGTTACGAC-3’) according to Chen and Yu 2000. A 473-bp product of the 16S rRNA gene was amplified via primers ANEM16SA (5’-CACTGACCGTGATAATG-TAGCGT-3’) and ANEM16SB (5’-CCCCATGGTAGCTTTTATTCG-3’) according to Geller and Walton 2001. Finally, a 721-bp product of the cytochrome oxidase III (COIII) gene was amplified via primers COIIIF (5’-CATTTAGTTGATCCTAGGCCTTGACC-3’) and COIIIR (5’-CAAACCACATCTA-CAAAATGCCAATATC-3’) according to Geller and Walton 2001. Finally, a 502-bp product of the 18S rRNA gene was amplified via primers 18S-3F (5’-GTTCGATTC-CGGAGAGGGA-3’) and 18S-5R (5’-CTTGGCAAATGCTITCGC-3’) according to Giribet et al. 1996. Annealing conditions of 55°C, 51.5°C, 51°C and 54°C were used for 12SrRNA, 16SrRNA, COIII, and 18S rRNA, respectively. Otherwise, all thermal protocols included the following steps: an initial 5 min denaturation at 94°C, then 1 min at 94°C,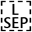1 min annealing step, and 1 min at 72°C, for 30 cycles, and a final 5 min extension at 72°C. Amplification products were sequenced directly using Sanger sequencing, via Laragen Inc., and submitted to GenBank (accession numbers xxxxx-xxxxx TBD, currently available at https://doi.org/10.5061/dryad.mkkwh70wt).

Newly generated DNA sequences for *Ostiactis pearseae* (and those for morphotype identified as Kadosactinidae sp.B. in this contribution) were combined and analyzed with the dataset by Gusmão et al. (2019) for each of the four markers (Table S2). Sequences for each marker were separately aligned in MAFFT v.7 (Katoh et al. 2013, 2017) using the following settings: Strategy, L-INS-I; Scoring matrix for nucleotide sequences, 200PAM/k = 2; Gap open penalty, 1.53; Offset value, 0.05. Alignments for each marker were analyzed separately and as a concatenated dataset (alignments available from the Dryad Digital Repository at https://doi.org/10.5061/dryad.mkkwh70wt). For each gene region, the best model of nucleotide substitution was chosen using the Akaike information criterion (AIC) on jModeltest2 (Guindon and Gascuel, 2003; Darriba et al. 2012) implemented on the CIPRES Portal (Miller et al., 2010). Maximum Likelihood (ML) analyses were performed using RAxML-NG v0.6.0 (Kozlov et al. 2018), using the appropriate model of nucleotide substitution for each gene partition (12S: GTR+I+G; 16S: TVM+G; COIII: TPM3uf+I+G; 18S: TIM2+I+G; 28S: GTR+I+G) in the combined alignment. The Majority Rule Criterion was used to assess clade support allowing bootstrapping to halt automatically (-autoMRE). All analyses were run with gaps treated as missing data.

### *Morphology and cnidae analysis of the anemone host* Ostiactis pearseae

Specimens were examined whole and dissected. Histological sections 5-10 μm thick were made from different body regions of two specimens using standard paraffin techniques and stained with Heidenhain Azan stain (Presnell and Schreibman, 1997). The distribution and size ranges of cnidae in the tissues was analyzed from six specimens using light DIC microscopy (1000x magnification, oil immersion). Twenty non-fired capsules of each cnida type (when possible) were photographed and measured at random. Cnidae size distribution offers information on the variability in capsule size for each type of nematocyst. We follow a nematocyst terminology that combines the classification of Weill (1934) modified by Carlgren (1940), thus differentiating ‘basitrichs’ from ‘*b*-mastigophores’ with that of Schmidt (1969, 1972) which captures the underlying variation seen in ‘rhabdoids’ (see Gusmão et al., 2018 for more details). We include photographs of each type of nematocyst for reliable comparison across terminologies and taxa (see Fautin, 1988). Higher-level classification for Actiniaria follows Rodríguez et al. (2014).

### Hybridization Chain Reaction-Fluorescent in-situ hybridization

Specimens for fluorescence in situ hybridization (FISH) microscopy were initially preserved in 4% sucrose-buffered paraformaldehyde (PFA) and kept at 4°C for 24-48 hours. These PFA-preserved specimens were then rinsed with 2× PBS, transferred to 70% ethanol, and stored at −20°C. Tissues were dissected and embedded in Steedman’s wax (1 part cetyl alcohol: 9 parts polyethylene glycol (400) distearate, mixed at 60°C). An ethanol: wax gradient of 3:1, 2:1 and 1:1, and 100% wax, was used to embed the samples (1 h each treatment at 37°C). Embedded samples were sectioned at ~3 μm thickness using a Leica RM2125 microtome and placed on Superfrost Plus slides. The protocol and all solutions used for HCR-FISH were as specified by Molecular Technologies, Inc., and closely followed Choi et al. 2014. Sections were dewaxed in three 100% ethanol rinses, allowed to dry, and equilibrated in hybridization buffer (Molecular Technologies; 30% formamide, 5× sodium chloride, sodium citrate (SSC = 750 mM NaCl, 75 mM sodium citrate), 9 mM citric acid (pH 6.0), 0.1% Tween 20, 50 μg/mL heparin, 1× Denhardt’s solution, 10% dextran sulfate), for 20 min at 37°C. Excess buffer was removed and sections were hybridized overnight in a humidification chamber at 37°C in hybridization buffer, to which was added a final concentration of 5 nM of an unlabeled DNA probe, designed to be an exact match to the *Ostiactis pearseae* SUP05 16S rRNA phylotype (Anem-SUP05, 5’-ACCATACTCTAGTTTGCCAG-3’), based on the probe, ‘BangT-642’, specific for the thiotrophic SUP05 symbiont in *Bathymodiolus* mussels (Duperron et al. 2005). A general bacterial probe set (Eub338-I-III) was also used as a positive control. These probes contained a specific sequence initiator tag (termed B1 and B3) that triggered the oligomerization of pairs of fluorescently-labeled DNA hairpins (i.e. the amplification step; Choi et al. 2014). The B1 initiator tag + linker (5’-GAGGAGGGCAGCAAACGG-GAAGAGTCTTCCTTTACG-ATATT-3’) was added to the 5’ end of the Anem-SUP05 probe. The B3 initiator tag + linker (5’-GTCCCTGCCTCTATATCTCCACTCAACTTT-AACCCG-ATATT-3’) was added to the 5’ end of each of three Eub338 probes I-III and to the Anem-SUP05 probe. In this case, tag B1 was paired with Alexa647-labelled hairpins, and tag B3 was paired with Alexa488-labelled hairpins.

Excess probe was removed by sequentially washing the slides for 15 min at 37°C in probe wash buffer (Molecular Technologies; 30% formamide, 5× SSC, 9 mM citric acid (pH 6.0), 0.1% Tween 20, 50 μg/mL heparin) to which 5× SSCT (750 mM NaCl, 75 mM sodium citrate, 0.1% Tween 20, pH 7) had been added to final concentrations (vol/vol) of 25%, 50%, and then 75%. This wash sequence was followed by two 15-min washes in 100% 5× SSCT at 37°C. Before amplification, 6 pmol of each hairpin, per reaction, was ‘snap cooled’ by heating to 95°C for 90 s, followed by 25°C for 30 min, in a thermocycler in separate PCR tubes. During this time, sections were equilibrated with amplification buffer (Molecular Technologies; 5× SSC, 0.1% Tween 20, 10% dextran sulfate) at room temperature for 30 min. For amplification, each ‘snap-cooled’ hairpin in a pair was added to 100 μl amplification buffer (for a final hairpin concentration of 60 nM for each amplifier hairpin), and then sections were incubated overnight (~18 h) at room temperature on a rocking platform protected from light. To remove unbound hairpin sequences, sections were washed twice in 5× SSCT for 15 min at room temperature, followed by two 30-min washes in 5× SSCT. Sections were rinsed with distilled water and counterstained with 4’6’-diamidino-2-phenylindole (DAPI, 5 mg/mL) for 1 min, rinsed again and mounted in Citifluor. Tissues were examined by epifluorescence microscopy using either a Nikon E80i epifluorescence microscope with a Nikon DS-Qi1Mc high-sensitivity monochrome digital camera or a Zeiss Elyra microscope with an ANDOR-iXon EMCCD camera.

### Transmission electron microscopy

Specimens for TEM and semi-thin sectioning were fixed in PFA and preserved in 50% EtOH. Before embedding, specimens were rehydrated, post-fixed with 1% OsO4 and subsequently dehydrated again in an ascending acetone series and embedded in araldite. 1 μm semi-thin sections were sectioned using a “Diatome Histo Jumbo” diamond knife on a Leica Ultracut S ultramicrotome and stained with toluidine blue (1% toluidine,1% sodium-tetraborate and 20% saccharose). Coverslips were mounted with araldite and sections were imaged with an Olympus microscope (BX-51) equipped with the dot.slide system (2.2 Olympus, Hamburg). Silver interference–colored sections (70 – 75 nm) were prepared using a “Diatome Ultra 45°” diamond knife. The sections were placed on Formvar-covered, single-slot copper grids and stained with 2% uranyl acetate and lead citrate in an automated TEM stainer (QG-3100, Boeckeler Instruments). Ultra-thin sections were examined using a Zeiss EM10 transmission electron microscope with digital imaging plates (DITABIS Digital Biomedical Imaging Systems, Germany).

## Declarations

Ethics approval and consent to participate

Not applicable

## Consent for publication

Not applicable

## Availability of Data and Materials

Longer partial length 16S rRNA and ITS sequences are available from GenBank under accession numbers XXXXXX-XXXXXX (TBD). The raw Illumina barcode sequence data and QIIME processed data are available in DataFile S1 from the Dryad Digital Repository (https://doi.org/10.5061/dryad.mkkwh70wt), along with representative sequences for the SUP05 OTUs. Alignments for the anemone 12S rRNA, 16S rRNA, 18S rRNA, and COIII for both *Ostiactis pearseae* and Kadosactinidae ‘sp.B’, used to generate Figure 3, are also available at https://doi.org/10.5061/dryad.mkkwh70wt. Animal images and specimens were vouchered for long-term archiving into the Benthic Invertebrate Collection at Scripps Institution of Oceanography (sioapps.ucsd.edu/collections/bi/).

## Competing interests - TBD

The authors declare that they have no competing interests.

## Authors’ Contributions

S.K.G. conducted DNA analysis, including 16S rRNA barcoding, host gene sequencing, and fluorescent microscope analyses, analyzed experimental data, wrote the manuscript with input from coauthors, and participated in the research expedition. C.M. conducted DNA analysis, including 16S rRNA barcoding. E.T. performed electron microscopy analyses and participated in the research expedition. D.F. performed the isotope analyses and reviewed the paper. G.W.R. interpreted the electron microscopy analyses, advised on the species identification, and participated in the research expedition. L.G. conducted phylogenetic analysis and advised on host identification. E.R. conducted host anemone analyses for species identification, including morphology, and wrote the manuscript. All authors contributed to data interpretation and editing of the paper.

## Acknowledgements

We thank the captain and crew of the R/V *Falkor, ROV SuBastian* pilots and technicians, as well as scientific participants of FK181031, chief scientists R. Zierenberg, D. Caress, and V. Orphan, as well as J. Paduan (MBARI) for Figure 1 maps and S. Connon for assistance with microbial community analysis.

## Supplemental Information - Goffredi et al.

**Table S1:** Number of total 16S rRNA amplicon reads, the Shannon Diversity index (H’) and the relative abundance (%) of the SUP05 group (based on 16S rRNA barcode amplification) associated with *Ostiactis pearseae*, water samples, and other Anthozoa from the Pescadero Basin vents.

| *Ostiactis pearseae* |  |  |  |  |
| --- | --- | --- | --- | --- |
| Auka vent field | Dive #-Sample # | # of reads | $H'$ index<sup>1</sup> | % SUP05 |
| Matterhorn | S0193-R2 | 11596 | 2.13 | 96.4 |
| E. of Diane's vent | S0193-A2 | 1852 | 2.79 | 70.9 |
| Z vent (top) | S0194-S2 | 2405 | 2.60 | 82.5 |
| S. of Z vent (small chimney) | S0200-R2w | 21299 | 2.67 | 87.0 |
| **JaichMaa 'ja'ag vent field** |  |  |  |  |
| Abuelita | S0197 | 16566 | 2.90 | 64.4 |
| Abuelita | S0197-CM2 | 4356 | 2.75 | 86.2 |
| Abuelita | S0197-CM3 | 6028 | 2.97 | 70.9 |
| Weey 'kual | S0199-S8 | 10220 | 2.61 | 69.4 |
| Water Samples |  |  |  |  |
| Auka vent field | Dive #-Sample # | # of reads | $H'$ index<sup>2</sup> | % SUP05 |
| E. of Diane's vent | S0193-N2 | 15754 | 4.38 | 6.7 |
| **JaichMaa 'ja'ag vent field** |  |  |  |  |
| Abuelita | S0197-N2 | 24802 | 4.91 | 6.0 |
| Weey 'kual | S0199-N1 | 10457 | 4.77 | 5.5 |
| Other Anthozoa |  |  |  |  |
| Auka vent field | Dive #-Sample # | # of reads | $H'$ index<sup>2</sup> | % SUP05 |
| Matterhorn | S0193-S4 | 1377 | 2.87 | 0.0 |
| S. of Z vent (small chimney) | S0200-R2r | 2805 | 3.14 | 0.3 |
| E. of Diane's vent | S0193-R3 | 40902 | 2.82 | 0.1 |
| Z vent (lower on structure) | S0194-R1 | 2858 | 3.64 | 0.4 |
| NW. of Z vent (diffuse flow) | S0194-R2 | 2759 | 3.19 | 0.3 |
| NW. of Z vent (diffuse flow) | S0194-S1 | 47970 | 2.46 | 0.0 |

<sup>1</sup> based on OTUs, defined as 99% similar based on 16S rRNA

**Table S2:**
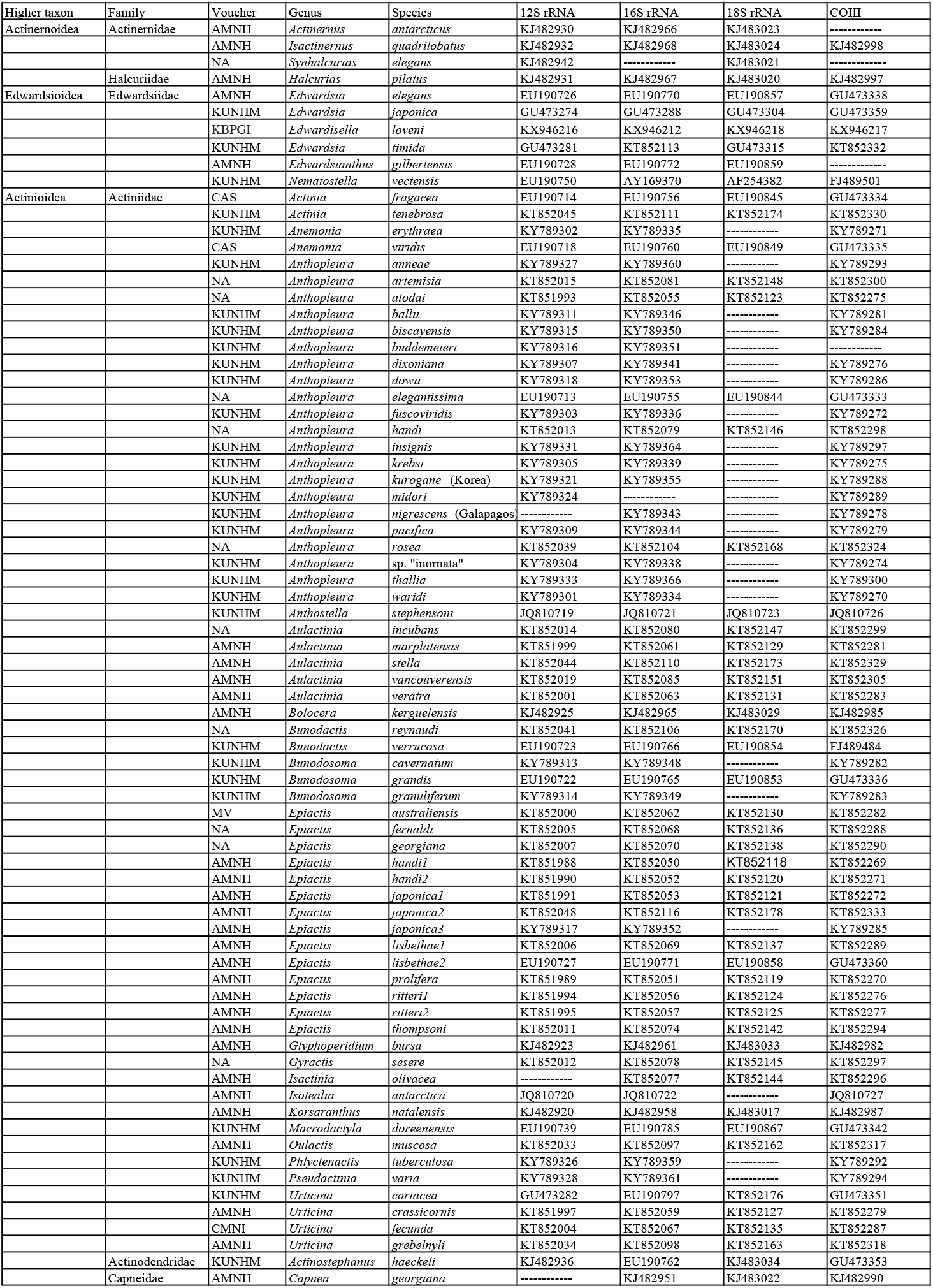
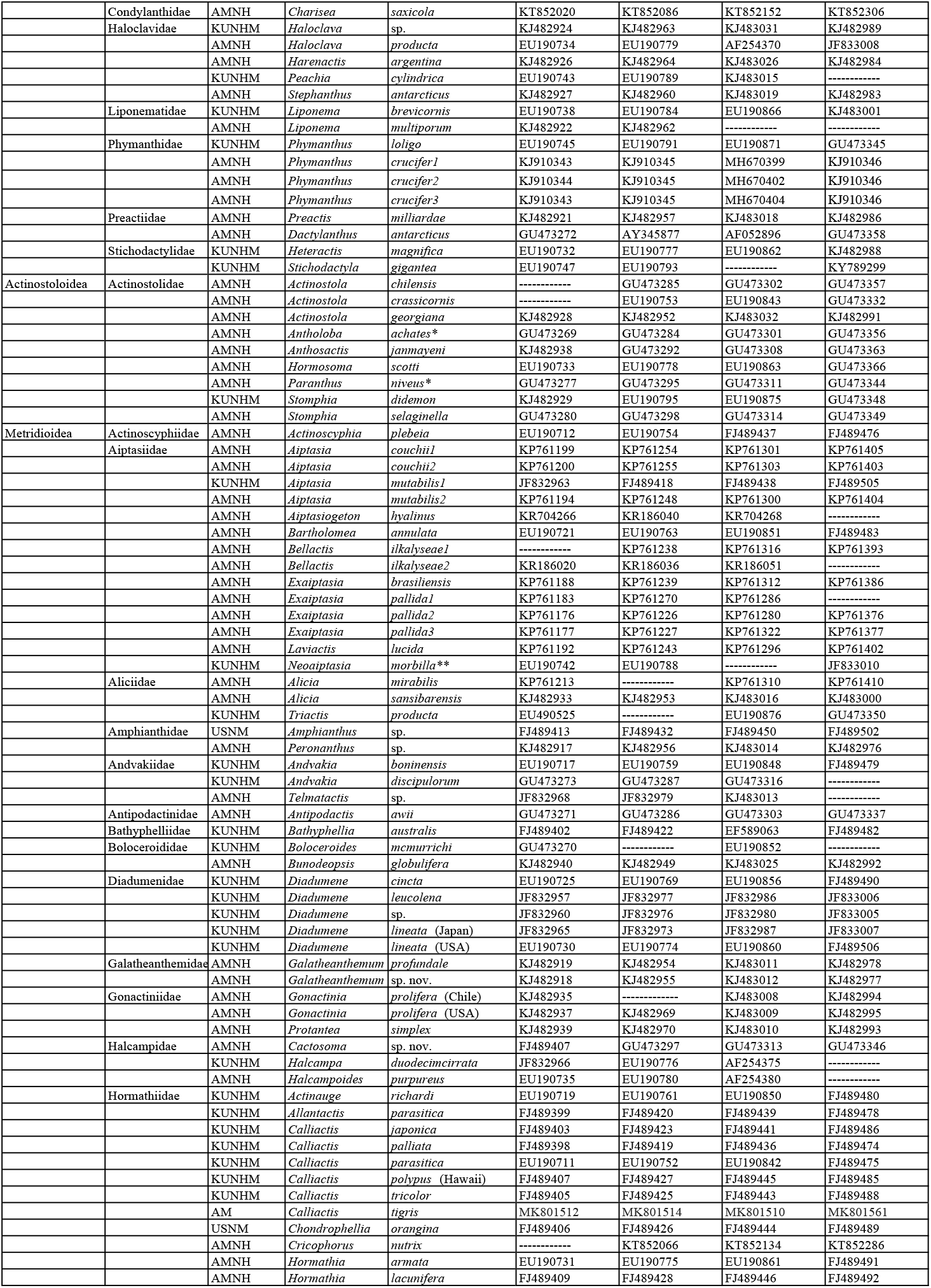
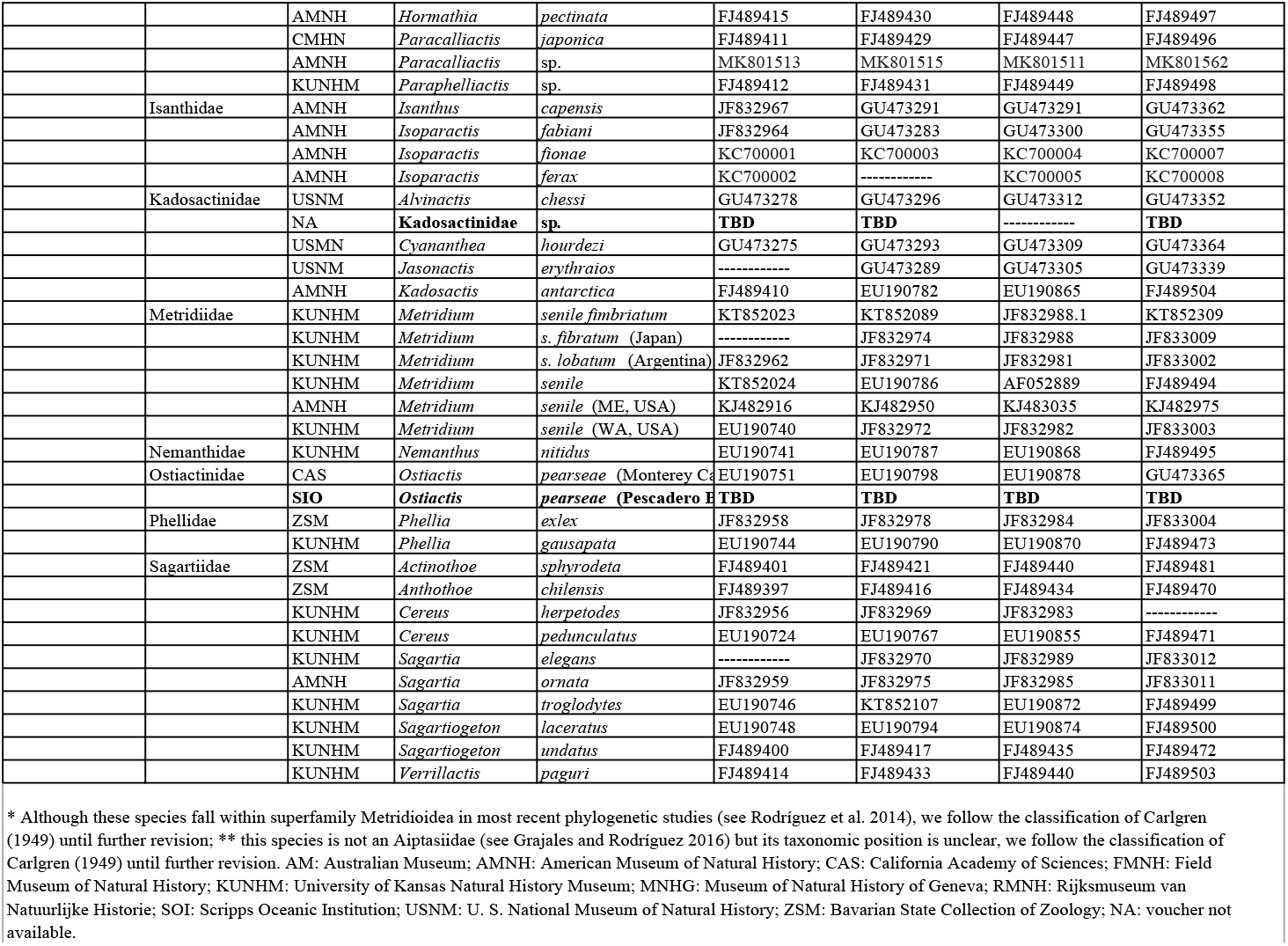
Taxa included in this study, with voucher location and GenBank accession numbers. Taxa are organized alphabetically within their family; new sequences indicated in bold.

**Fig. S1.**
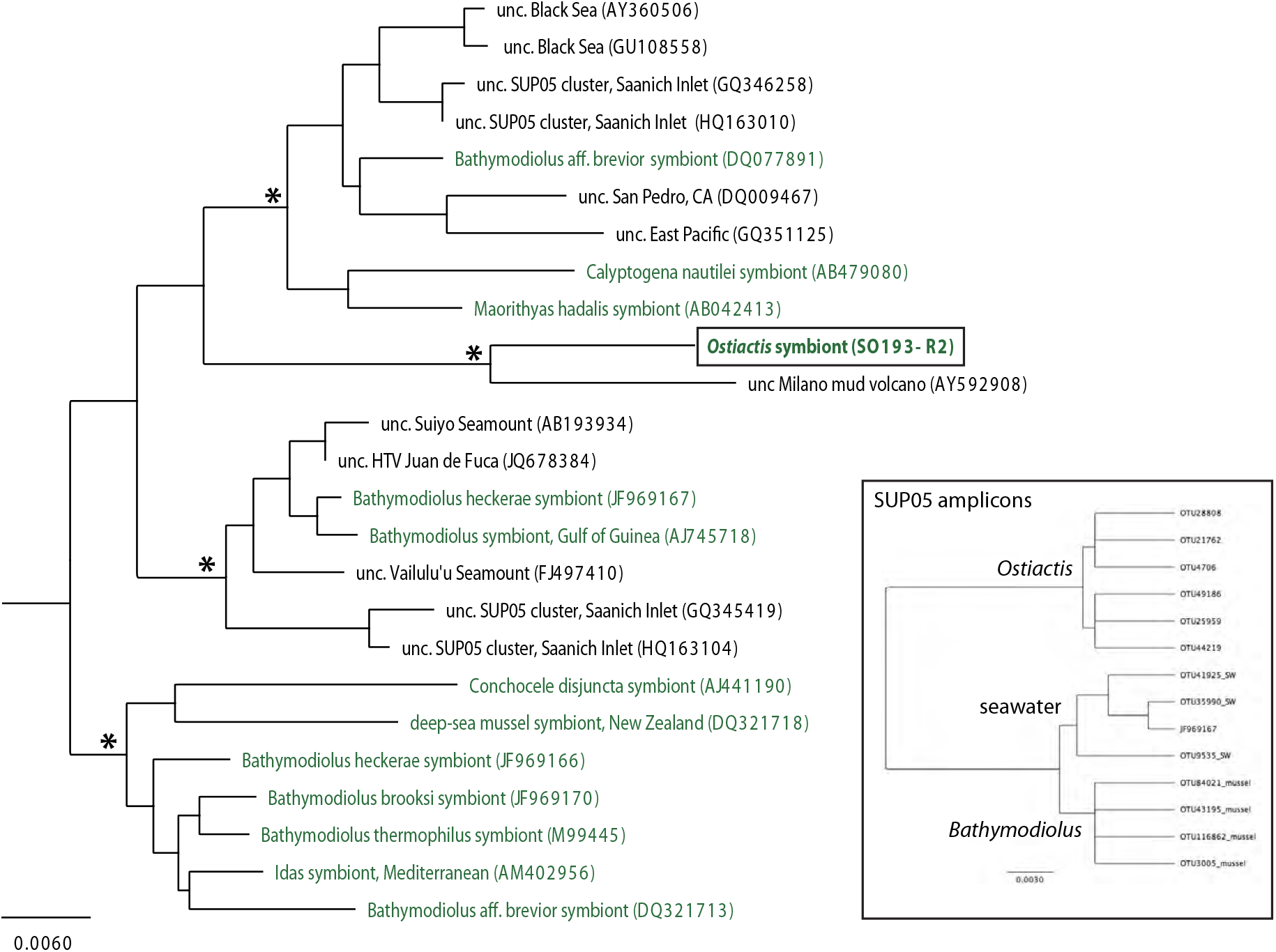
Phylogenetic relationships of the SUP05 group, based on 16S rRNA. **A.** SUP05 cluster, based on 16S rRNA. Taxa shown in green are known symbionts of marine invertebrates. * > 70% support (using the Jukes Kantor model). Additional taxa were included according to Petersen et al. 2012; Glaubitz et al. 2013; Shah et al 2019. **Inset**. Shows SUP05 amplicons recovered from *Ostiactis pearseae*, surrounding seawater samples, and *Bathymodiolus* mussels from the Costa Rica margin Jaco Scar seep sites (SG, unpublished).

**Fig. S2.**
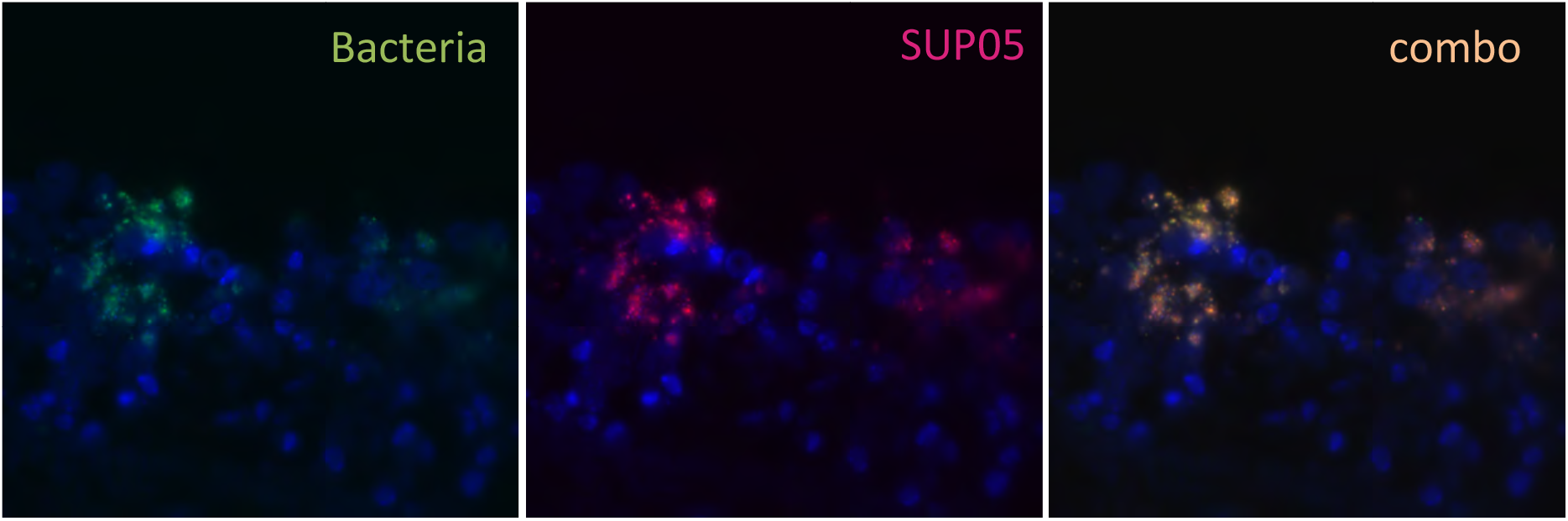
Fluorescence Microscopy of the tentacles of *Ostiactis pearseae.* Fluorescent *in situ* signal amplification via hybridization chain reaction-FISH (HCR-FISH) microscopy of *Ostiactis pearseae* tentacles using **A.** a general bacterial probe set Eub338 I-III, **B.** the specific Anem_SUP05 probe, and **C.** an overlay of the two showing near complete overlap. Scale is 10 μm.

Available from the Dryad Digital Repository (https://doi.org/10.5061/dryad.mkkwh70wt)

**Supp Video**: Sea anemones were collected by ROV manipulator or suction sampler mounted on ROV *SuBastian.*

## DataFile S1

The raw Illumina barcode sequence data and QIIME processed data, including representative sequences for all OTUs, as well as representative sequences for the SUP05 OTUs specifically.

